# HRV-GUI: A MATLAB Graphical User Interface for Heart Rate Variability Analysis and Validation Using Human, Rodent, and Clinical Diabetic Gastroparesis Data

**DOI:** 10.64898/2026.09.01.748623

**Authors:** M. Khawar Ali, Jiande D. Z. Chen

## Abstract

**Background and Objective:** Heart rate variability (HRV) analysis provides a non-invasive method for quantifying autonomic modulation from electrocardiographic recordings. However, practical HRV analysis often depends on fragmented workflows, limited signal-quality review, and software tools optimized for either human or preclinical recordings, but not both. This study developed and evaluated HRV-GUI, a MATLAB-based graphical interface for electrocardiogram (ECG)-derived HRV analysis in translational biomedical research.

**Methods:** The HRV-GUI integrates electrocardiographic and RR interval loading, human and rat analysis modes, preprocessing, segment selection, automated R-peak detection, manual peak correction, RR interval generation, multi-domain HRV computation, diagnostic visualization, result export, and session saving/loading. The software was evaluated using deterministic synthetic RR interval datasets, baseline recordings from healthy human controls and healthy rats, and a clinical use-case comparison between healthy controls and patients with diabetic gastroparesis.

**Results:** The HRV-GUI produced expected outputs in synthetic RR validation tests, including constant RR sequences, alternating RR sequences, outlier-containing RR sequences, and low-frequency- or high-frequency-dominant sinusoidal RR modulation. The software generated physiologically plausible HRV profiles in both human and rat recordings. In the clinical use-case analysis, patients with diabetic gastroparesis showed higher heart rate and sympathetic index, together with lower respiratory sinus arrhythmia, absolute low- and high-frequency spectral power, standard deviation of normal-to-normal intervals (SDNN), root mean square of successive differences (RMSSD), percentage of successive RR intervals differing by more than 50 ms (pNN50), Poincaré short-term variability (SD1), and Poincaré long-term variability (SD2) compared with healthy controls.

**Conclusions:** HRV-GUI provides an integrated biomedical software workflow for ECG-derived HRV analysis. The validation results support its use for controlled RR testing, human and rodent ECG recordings, and clinical autonomic assessment in diabetic gastroparesis.

## 1. Introduction

Heart rate variability (HRV) describes beat-to-beat fluctuation in sinus rhythm and is widely used as a non-invasive marker of autonomic nervous system regulation. HRV reflects dynamic modulation of cardiac rhythm by sympathetic and parasympathetic inputs and has been applied across cardiovascular, neurological, gastrointestinal, metabolic, and behavioral research (Task Force of the European Society of Cardiology and the North American Society of Pacing and Electrophysiology, 1996; Thayer and Sternberg, 2006; Shaffer and Ginsberg, 2017).

In gastrointestinal research, autonomic dysfunction has been implicated in disorders of gut-brain interaction and motility disorders, including irritable bowel syndrome, gastroparesis, chronic constipation, and gastrointestinal dysmotility (Mazurak et al., 2012; Ali et al., 2021; Liu et al., 2022; Ali and Chen, 2023). ECG-derived HRV provides a practical approach for quantifying autonomic regulation in these settings because RR interval series can be analyzed using established time-domain, frequency-domain, and nonlinear metrics (Task Force, 1996; Shaffer and Ginsberg, 2017). Previous work has also shown that HRV parameters can quantify autonomic responses to brief rectal distention in patients with irritable bowel syndrome, supporting the use of HRV for gastrointestinal autonomic assessment (Ali et al., 2023).

Despite the broad use of HRV, practical barriers remain in routine biomedical research workflows. Many laboratories rely on custom scripts, commercial packages, or fragmented analysis pipelines that may be difficult to reproduce, difficult for non-programmers to use, or not designed for both human and rodent recordings. Existing HRV and biomedical signal-processing tools provide important capabilities for HRV computation, artifact handling, and physiological signal analysis, but they may not fully address the combined needs of interactive ECG review, manual R-peak correction, RR interval inspection, and dual human/rodent processing within a single laboratory-facing interface (Tarvainen et al., 2014; Garcia, 2009; Kaufmann et al., 2011; Vidaurre et al., 2011). Open physiological signal repositories and software resources such as PhysioNet and PhysioToolkit have also supported reproducible biomedical signal analysis and method development (Goldberger et al., 2000). For translational autonomic research, an HRV tool should therefore do more than compute formulas. It should support signal inspection, artifact handling, robust R-peak detection, manual peak correction, RR interval review, diagnostic visualization, and structured export of results for downstream statistical analysis.

These requirements are particularly important for laboratories that analyze both clinical human ECG recordings and preclinical rodent ECG recordings. Human and rodent HRV recordings differ substantially in heart rate, RR interval scale, expected peak spacing, respiratory rhythm, spectral frequency ranges, and interpretation of some HRV parameters (Thireau et al., 2008; Baudrie et al., 2007). Therefore, a single interface that supports both species while preserving species-specific analytical settings can improve workflow consistency across translational studies.

The HRV-GUI was developed to address these needs. It is a MATLAB-based graphical user interface that integrates ECG loading, preprocessing, segment selection, R-peak detection and editing, RR interval generation, multi-domain HRV computation, diagnostic visualization, data export, and session management in a single environment. The purpose of this manuscript is to describe the software, summarize the ECG acquisition procedures used in the demonstration datasets, and present a multi-layer validation framework using deterministic synthetic RR datasets, cross-species baseline feasibility testing, and a clinical human use-case comparison between healthy controls and patients with diabetic gastroparesis.

## 2. Software description

### 2.1 Architecture and Design

The HRV-GUI was implemented in MATLAB as a graphical research interface for ECG-derived HRV analysis. The software allows users to analyze either raw ECG waveforms or pre-generated RR interval files and includes species-specific processing options for human and rat recordings.

The software supports ECG or RR interval loading, species selection, preprocessing, segment selection, automated and manual R-peak handling, RR interval generation, HRV computation, diagnostic visualization, result export, and session management.

The workflow shown in Figure 1 represents the standard analysis sequence used in this manuscript. For ECG-based analysis, the user launches HRV-GUI, selects the study type as human or rat, and loads a WAV-format ECG file. Study-type selection initializes species-appropriate peak-detection defaults and frequency-domain settings. The ECG waveform is then reviewed visually. During this step, the user may zoom, scroll, select a clean analysis segment, delete noisy segments, invert the ECG polarity if R-peaks are negative, or undo the most recent editing operation before peak detection.

**Figure 1.**
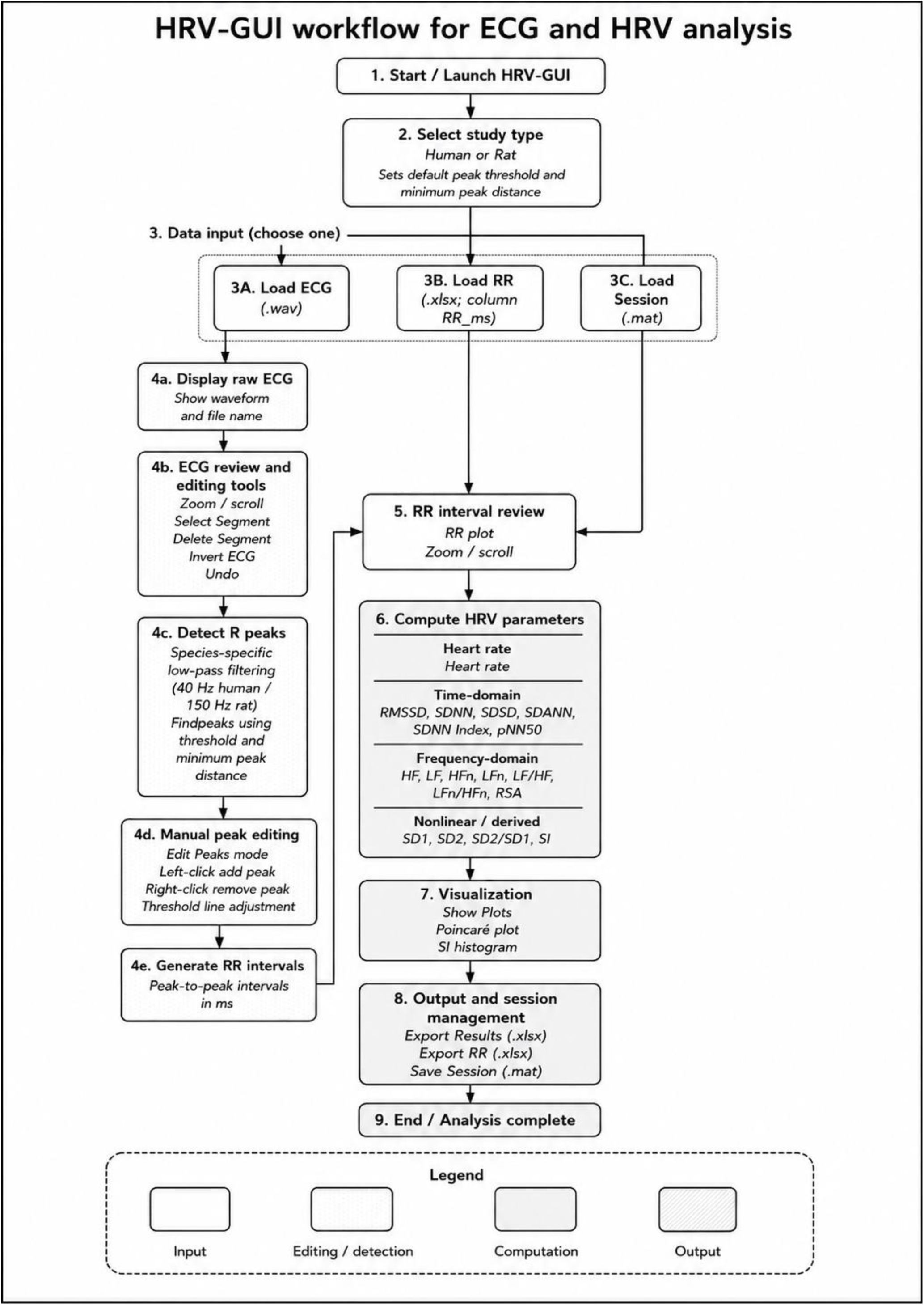
Conceptual HRV-GUI workflow. The software supports ECG or RR loading, species selection, preprocessing and segment selection, automated and manual R-peak handling, RR interval generation, HRV calculation, visualization, export, and session management.

After ECG review, the HRV-GUI applies species-specific preprocessing for R-peak detection and identifies candidate R-peaks using adjustable threshold and minimum-peak-distance criteria. Because automated peak detection may fail in the presence of noise, motion artifact, abnormal waveform morphology, or high-rate rodent ECG signals, the software includes an interactive Edit Peaks mode. In this mode, users can manually add missed peaks or remove false detections. Left-clicking adds a peak near the nearest local maximum, whereas right-clicking removes the nearest detected peak. This manual correction step is important because even a small number of missed or falsely detected R-peaks can substantially alter the RR interval series and downstream HRV indices.

Once R-peak locations are finalized, HRV-GUI generates the RR interval series as consecutive peak-to-peak differences in milliseconds. Alternatively, users can bypass ECG-based peak detection by loading a precomputed RR interval file containing a column named RR_ms. The software also supports MAT-session reloading, allowing users to restore the ECG signal, detected peaks, RR intervals, HRV results, study type, peak threshold, peak distance, and sampling frequency from a previously saved analysis session. After RR interval review, HRV-GUI computes HRV parameters, generates diagnostic plots, exports HRV and RR outputs to Excel, and allows the complete analysis session to be saved for reproducibility.

### 2.2 Main interface and user controls

The main interface is organized around two signal-display panels and a results table. The upper panel displays the ECG waveform with detected R-peaks, the middle panel displays the generated or loaded RR interval series, and the lower table displays the calculated heart rate and HRV outputs. The right-side control panel contains the study-type selector, peak-detection settings, signal-processing controls, analysis workflow buttons, export functions, and session-management options.

The upper ECG panel is used for waveform visualization and R-peak inspection. It can display the raw or processed ECG signal, with detected R-peaks overlaid as markers and a visible threshold line during peak detection. This allows the user to assess signal quality, inspect peak-detection performance, identify noisy or artifact-contaminated regions, and confirm whether the selected segment is suitable for HRV analysis. The middle RR panel displays the RR interval series after peak detection or direct RR-file loading, allowing the user to identify abrupt deviations, rhythm irregularities, or potential missed or false R-peaks before HRV computation. The lower results table provides immediate review of the calculated heart rate and HRV parameters before export.

The GUI controls are arranged to follow the intended analysis sequence: data loading and study-type selection, signal review and editing, R-peak detection, RR interval generation, HRV computation, diagnostic visualization, export, session management, and exit. This structure helps ensure that users review the ECG waveform and RR interval series before calculating final HRV outputs. The controls listed in Table 1 correspond directly to the workflow shown in Figure 1 and the interface layout shown in Figure 2. Figure 3 shows a representative HRV-GUI analysis session in human mode, including the preprocessed ECG signal with detected R-peaks, the generated RR interval trace, and the calculated HRV output table.

**Table 1.**
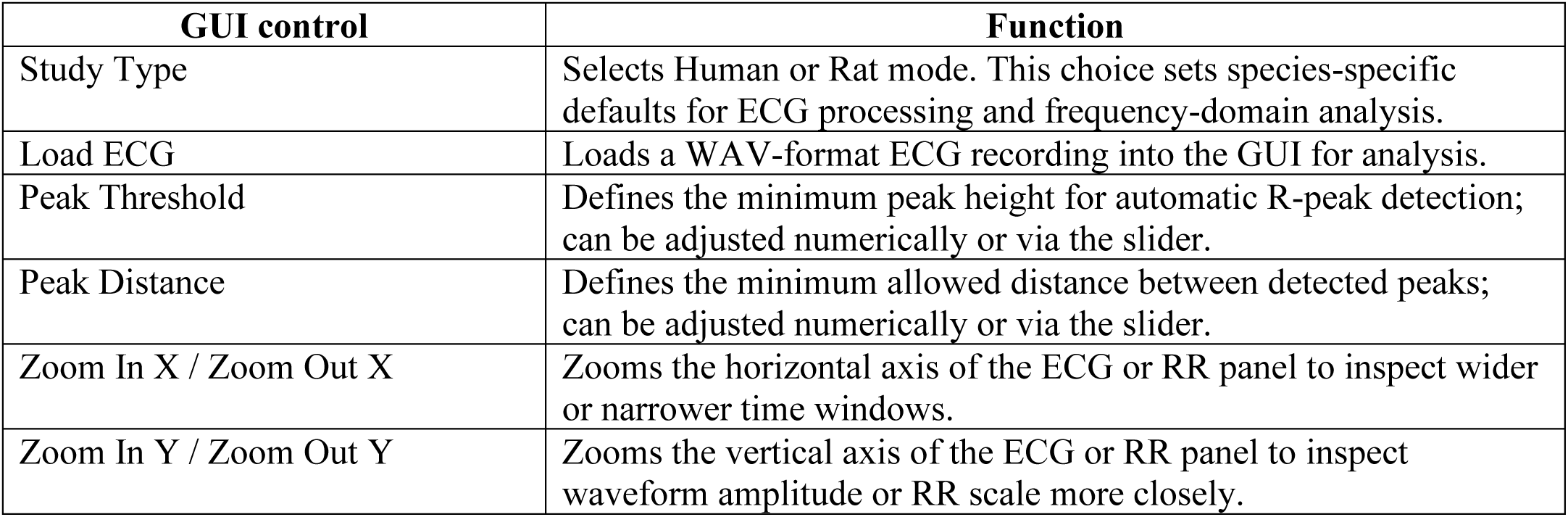
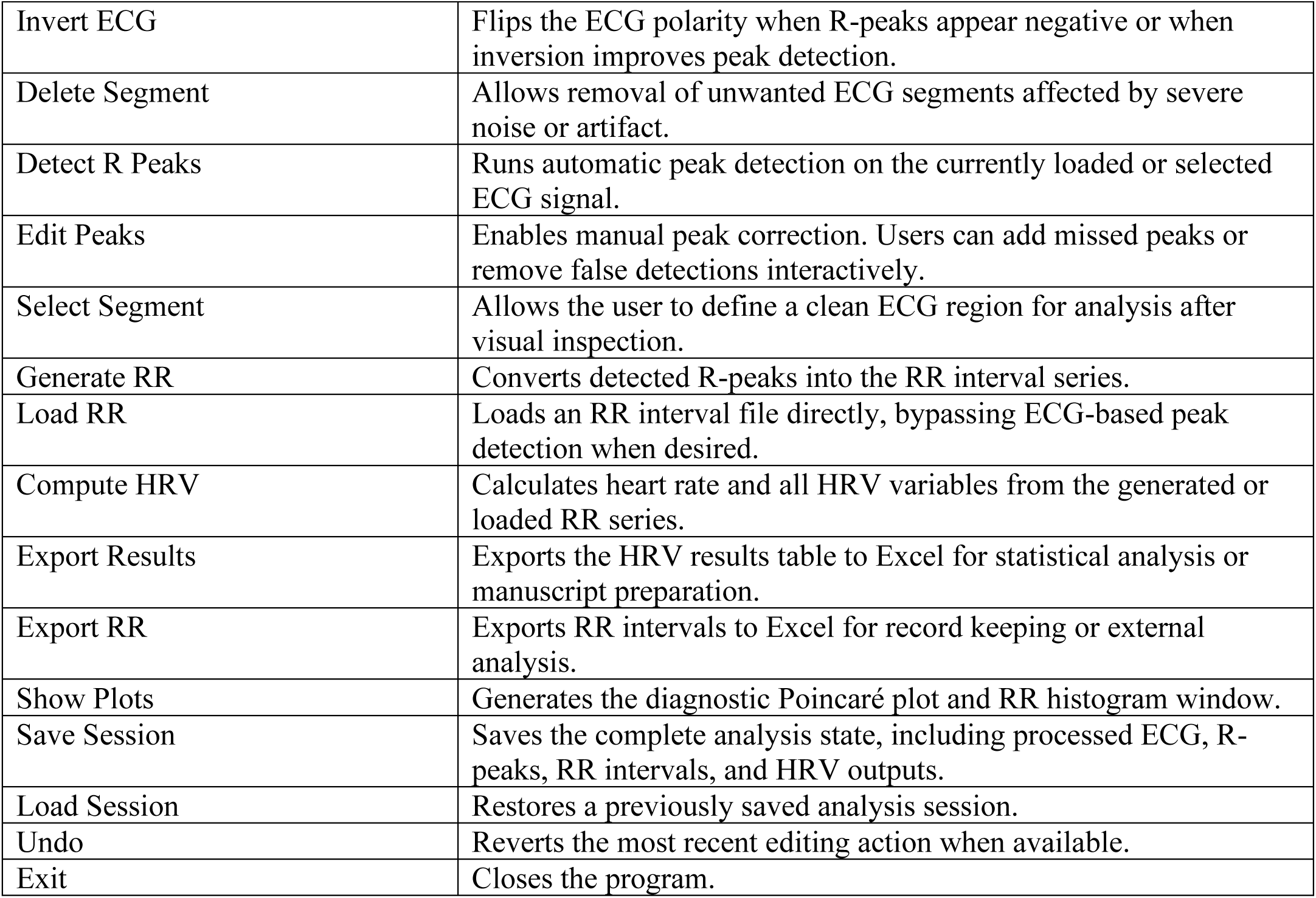
HRV-GUI user controls and their primary functions.

**Figure 2.**
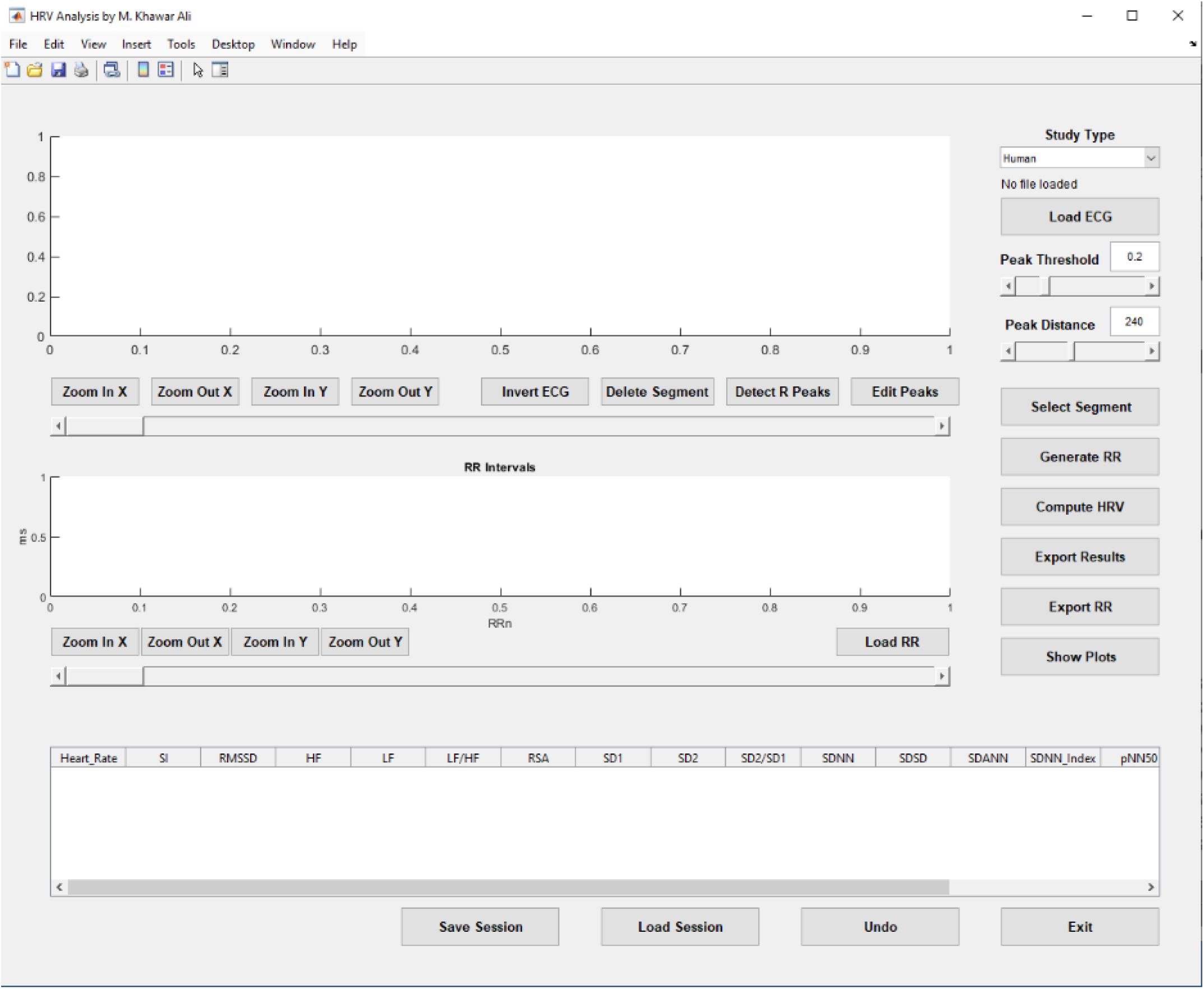
HRV-GUI main interface. (Top Panel) ECG display panel with zoom/scroll controls and signal processing buttons (Invert ECG, Delete Segment, Detect R Peaks, Edit Peaks). (Middle Panel) RR interval visualization panel with independent zoom and scroll controls. (Bottom Panel) Scrollable HRV results table displaying all 17 computed HRV parameters and heart rate. (Right Panel) Right-side control panel showing Study Type selector (Human/Rat), Load ECG button, Peak Threshold and Peak Distance controls with real-time sliders, and analysis workflow buttons (Select Segment, Generate RR, Compute HRV, Export Results, Export RR, Show Plots). (Bottom) Session management controls (Save Session, Load Session, Undo, Exit). Peak Threshold default = 0.2 mV; Peak Distance default = 240 samples (Human mode).

**Figure 3.**
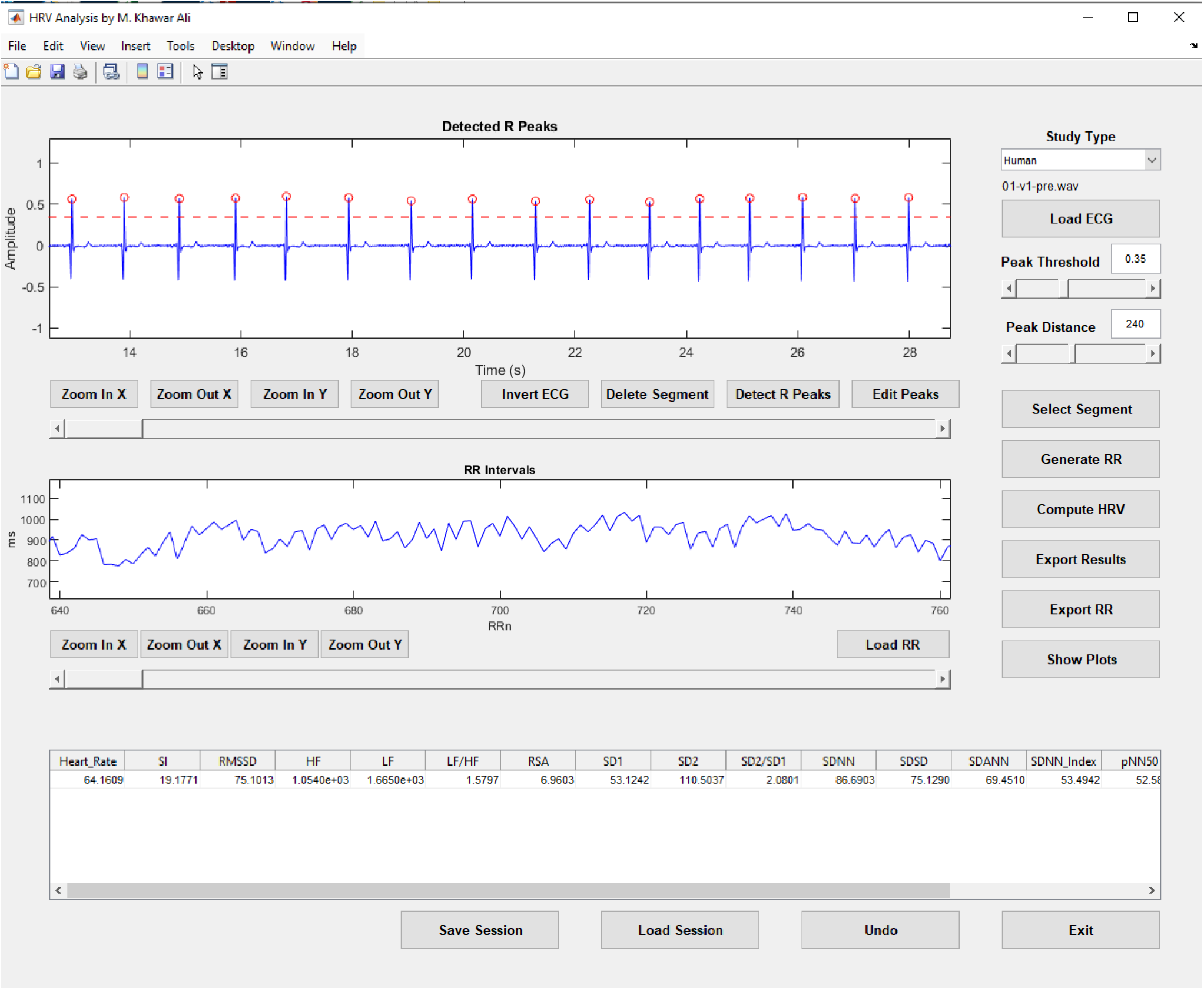
Representative HRV-GUI interface in human mode. The GUI displays the preprocessed ECG with detected R-peaks, the RR interval trace, and the calculated HRV values in the lower results table.

In routine ECG-based analysis, the user proceeds from study-type selection and ECG loading to signal inspection, segment selection, peak detection, manual peak correction when needed, RR interval generation, HRV computation, diagnostic visualization, and export. The interface also supports flexible non-linear workflows, such as loading precomputed RR interval files, reloading a previously saved session, correcting peaks before regenerating RR intervals, or re-exporting results after review. This design provides a structured but flexible workflow for HRV analysis in both human and rat recordings.

### 2.3 Signal loading, preprocessing, and R-peak handling

ECG recordings are loaded into HRV-GUI in WAV format, which is compatible with the direct output of the UFI amplifier used in our laboratory. After loading the file, the user specifies the study type as either human or rat. This selection is important because human and rodent recordings differ substantially in expected heart rate, RR interval scale, R-peak spacing, waveform characteristics, and frequency-domain HRV settings.

After ECG loading, the waveform can be visually inspected before analysis. The preprocessing workflow is designed to improve R-peak detection while preserving the physiologic timing of the R-wave. The software allows the user to invert ECG polarity when R-peaks appear negative, select a clean analysis segment, and remove severely noisy or artifact-contaminated regions when needed. These steps allow the user to focus HRV computation on an artifact-minimized ECG segment.

Automatic R-peak detection is controlled by adjustable peak-threshold and minimum-peak-distance settings. The threshold defines the minimum peak amplitude required for detection, whereas the peak-distance setting defines the minimum allowable interval between adjacent detected peaks. These settings can be adjusted according to species, signal amplitude, and recording quality. Because fully automated peak detection may not be reliable in all research recordings, HRV-GUI also includes interactive manual peak editing. Users can add missed R-peaks or remove false detections before RR interval generation. This feature is particularly useful in motion-contaminated human recordings and high-rate rat ECG recordings, where even a small number of missed or falsely detected peaks can substantially alter the RR interval series and downstream HRV parameters.

### 2.4 RR interval generation, HRV outputs, and diagnostic plots

Once the ECG peaks are finalized, the software generates the RR interval series and computes heart rate and HRV parameters spanning time-domain, frequency-domain, nonlinear, and Poincaré domains. These domains follow established HRV measurement standards and commonly used HRV metric definitions (Task Force, 1996; Shaffer and Ginsberg, 2017). The variables generated by HRV-GUI are listed in Table 2, together with their units and implementation descriptions. Poincaré-derived SD1 and SD2 were interpreted according to established geometric descriptions of short- and long-axis RR interval variability (Brennan et al., 2001), while the SI was implemented according to the Baevsky stress-index formulation (Baevsky and Berseneva, 2008). Human frequency bands followed conventional short-term HRV standards (Task Force, 1996), whereas rat frequency bands were selected based on published rodent HRV guidance (Thireau et al., 2008; Baudrie et al., 2007).

**Table 2.**
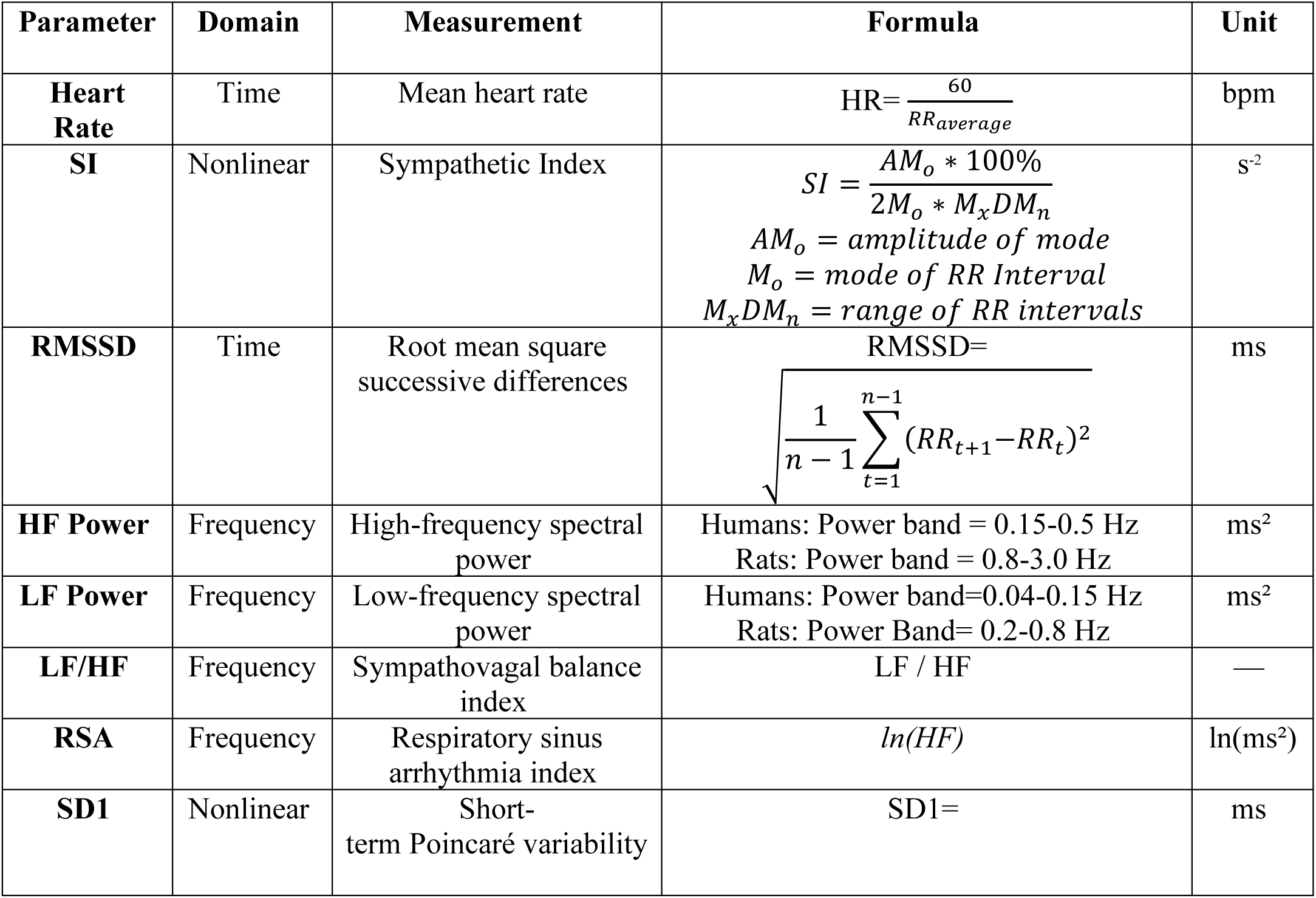
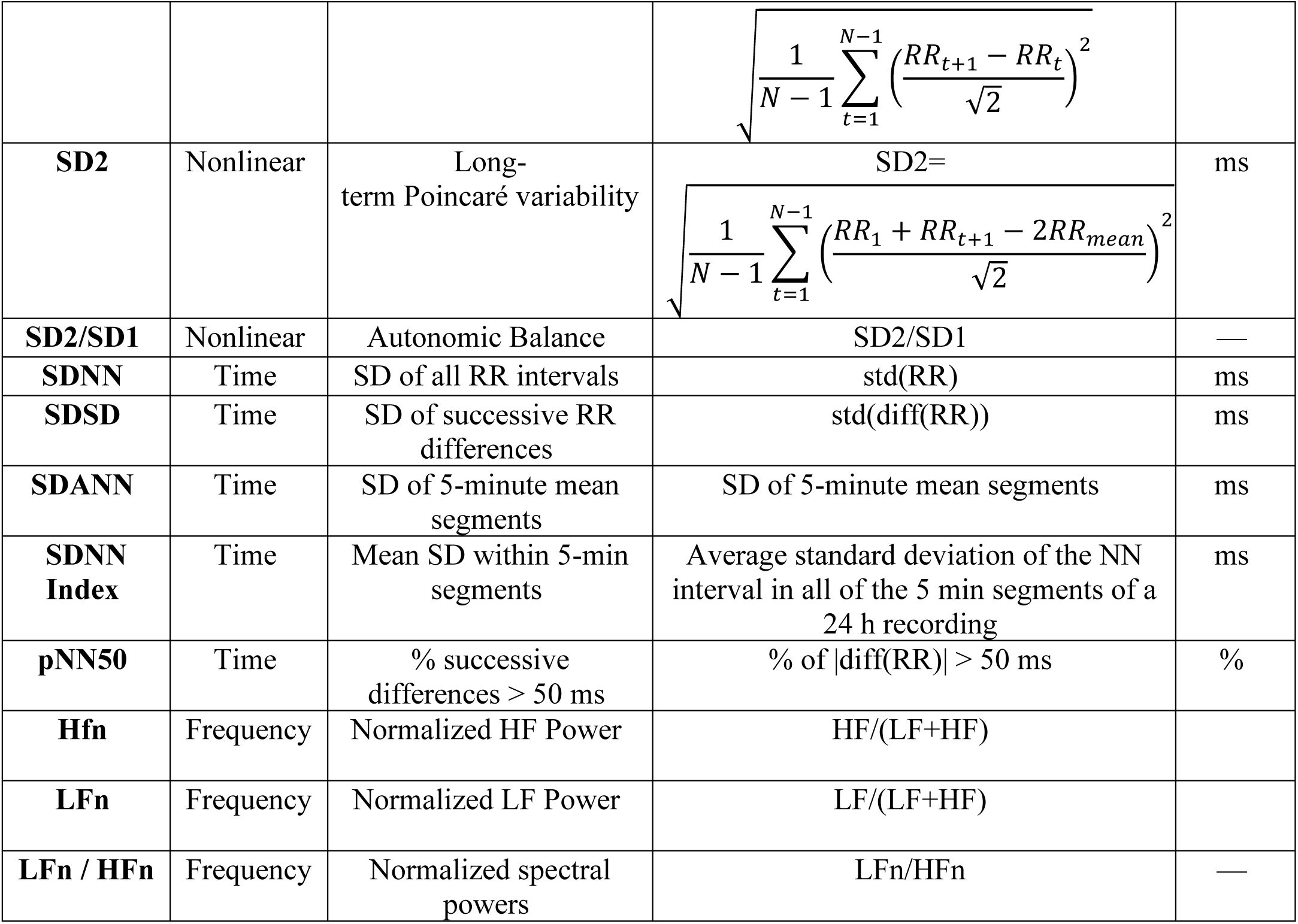
HRV variables reported by HRV-GUI in this manuscript. Human recordings used human HRV frequency bands, whereas rat recordings used rat-specific frequency bands.

In addition to the numerical HRV output table, HRV-GUI generates diagnostic visualizations to help users review RR interval structure and Poincaré-derived variability. Representative human diagnostic output is shown in Figure 4, including the Poincaré plot and RR interval histogram. A representative rat-mode output is shown in Figure 5, demonstrating ECG peak detection, RR interval generation, HRV calculation, and rodent-specific diagnostic visualization using rat-adjusted frequency-domain settings.

**Figure 4.**
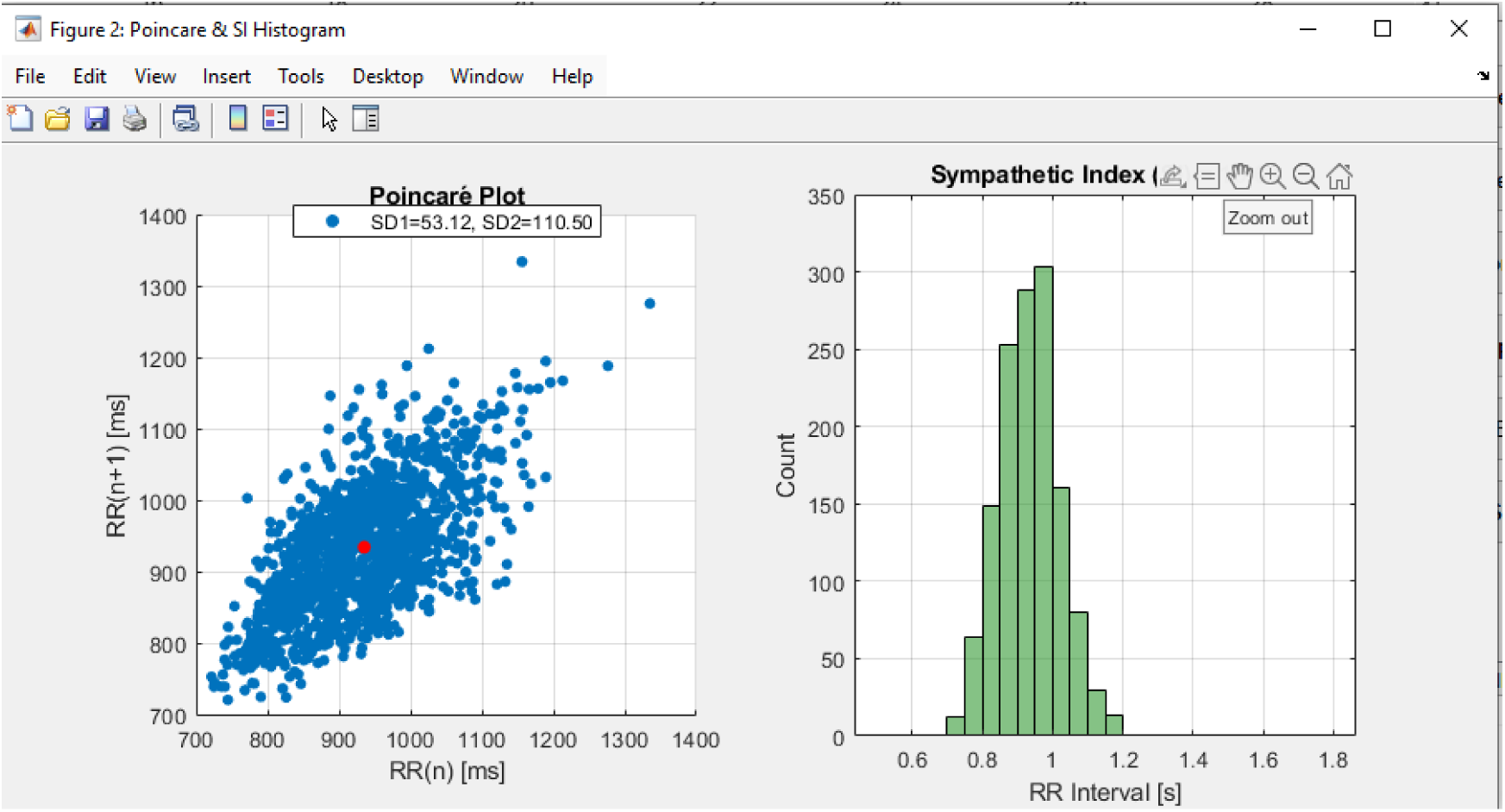
Representative human diagnostic output generated by HRV-GUI, showing the Poincaré plot and RR interval histogram. The Poincaré panel reports SD1 and SD2, while the histogram visualizes RR interval distribution used in stress-index-related interpretation.

**Figure 5.**
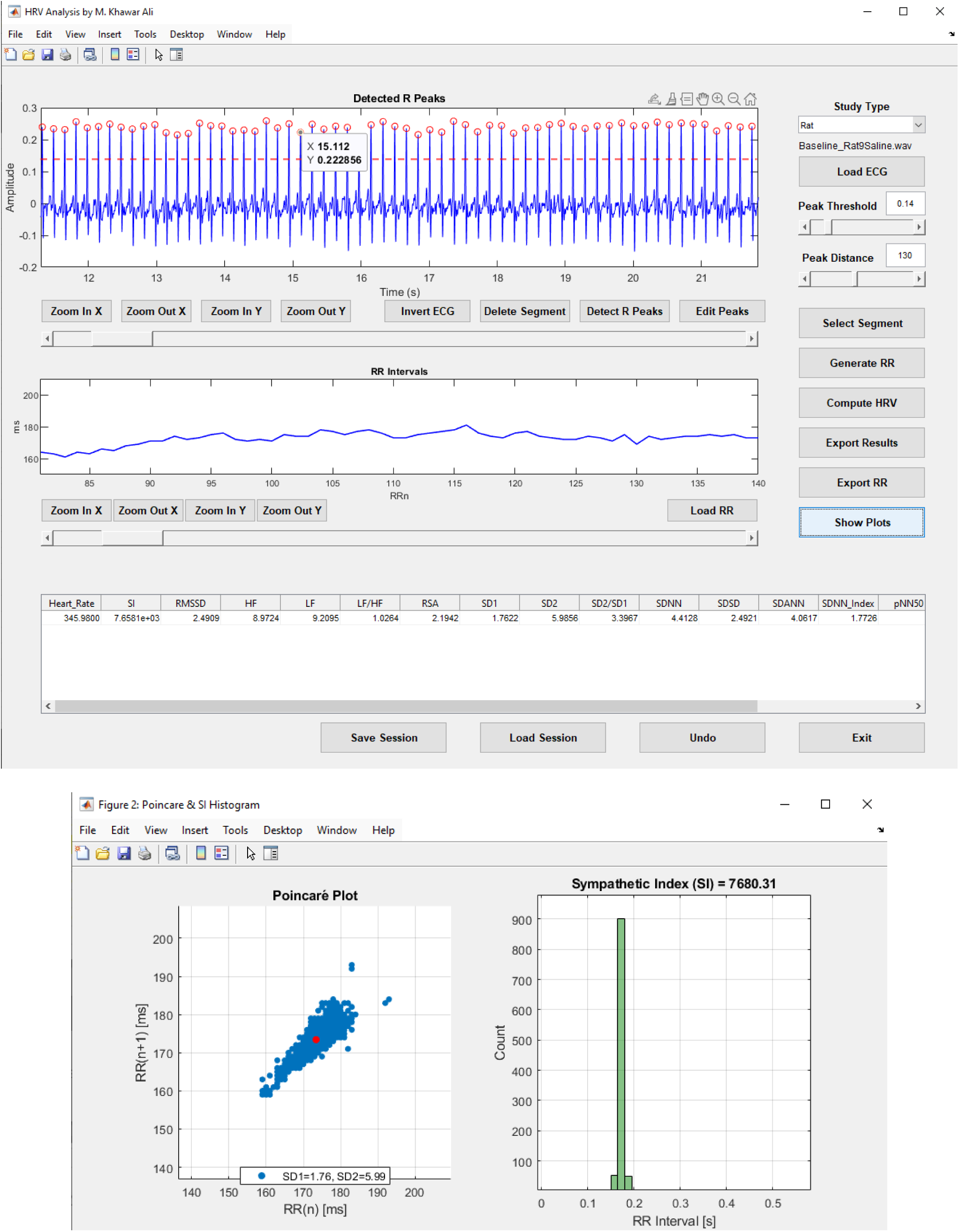
Representative HRV-GUI output in rat mode. The upper panel shows a representative rat ECG segment with detected R-peaks, generated RR intervals, and calculated HRV values. The lower panel shows the rat Poincaré plot and RR histogram. The tighter Poincaré distribution compared with the human example reflects the smaller absolute RR variability typical of rodent recordings. Rodent-adjusted frequency bands were applied automatically.

## 3. ECG recording procedures

### 3.1. Human ECG recordings

Human ECG recordings were obtained in a seated position under a standardized condition. Recordings from patients with diabetic gastroparesis were conducted in the GI motility clinic at Michigan Medicine. Healthy control recordings were conducted in our research office under controlled environmental conditions. For both groups, participants were seated comfortably and were instructed to remain still, avoid talking, and breathe regularly during acquisition. Lighting, room temperature, seating position, and recording posture were kept as consistent as possible to minimize environmental and procedural variation, because HRV can be influenced by posture, arousal, respiration, temperature, and other recording conditions. The recordings were made in the fasting state.

Human ECG signals were acquired using the UFI Universal Fetrode Amplifier (model 2283 FTI/2283FT; UFI, Morro Bay, CA, USA). Three surface electrodes were used for ECG acquisition, including positive, negative, and reference electrodes. Recordings were approximately 30 minutes in duration.

### 3.2. Rat ECG recordings

Rodent ECG recording was made via chronically implanted electrodes. Under inhaled isoflurane anesthesia at 1.5-2.0%, the chest hair was shaved and the skin was disinfected. Three small chest skin incisions, approximately 2 mm each, were made, and three cardiac pacing wire electrodes (Medtronic, Minneapolis, MN, USA) were implanted subcutaneously in the left chest, right chest, and left cardiac-apex region. The lead wires were tunneled subcutaneously and externalized percutaneously at the neck using a procedure adapted from Wang et al. (2019).

After surgery, carprofen (5 mg/kg) and enrofloxacin (5 mg/kg) were administered subcutaneously once daily for 3 days beginning on the day of surgery to provide postoperative analgesia and support recovery. Rats were housed individually to protect the externalized electrode wires from chewing by cage mates. Experiments were performed after a 7-day recovery period following electrode implantation.

After recovery, rats were fasted overnight before the experiment. On the day of the experiment, each rat was placed in a transparent restrainer, and the ECG was recorded for 30 minutes using a UFI Universal Fetrode Amplifier (model 2283 FTI/2283FT; UFI, Morro Bay, CA, USA). The resulting ECG recordings were analyzed in the HRV-GUI using rat-specific peak-detection and frequency-domain settings.

## 4. Validation framework and statistical analysis

The HRV-GUI was evaluated using three complementary validation levels: (1) deterministic synthetic RR interval datasets to test calculation logic and edge-case behavior, (2) cross-species baseline feasibility testing in healthy human controls and healthy rats to determine whether the same software workflow could process recordings from both species using species-specific settings, and (3) a clinical use-case comparison between healthy controls and patients with diabetic gastroparesis to demonstrate biomedical applicability in real human ECG recordings.

This validation structure was designed to test different components of the software workflow. Synthetic RR files evaluated the HRV calculation engine independently of ECG signal quality and R-peak detection. Cross-species baseline analysis evaluated whether HRV-GUI could generate physiologically plausible outputs from real human and rat ECG recordings. The HC-DGP comparison evaluated whether the complete workflow, including ECG processing, RR interval generation, HRV computation, diagnostic visualization, and statistical export, could generate clinically interpretable group differences in a representative human use case.

For the synthetic RR validation, deterministic RR interval files were generated with known expected behavior, including constant RR sequences, alternating RR sequences, outlier-containing RR sequences, and LF- or HF-dominant sinusoidal RR modulation. These files were loaded directly into HRV-GUI using the RR-loading function so that the calculation engine could be evaluated independently of ECG peak detection.

For the cross-species feasibility analysis, healthy human and healthy rat baseline recordings were summarized descriptively. Human and rat values were presented in separate tables because the two species differ substantially in heart rate, RR interval scale, autonomic physiology, respiratory rhythm, and frequency-domain definitions. No formal inferential statistics were performed between human and rat groups. Descriptive statistics included mean ± SD, median with interquartile range, and observed range.

Normality was assessed separately for each HRV variable within each group. When both groups were approximately normally distributed, between-group comparisons were performed using Welch’s unequal-variance two-sample t-test. When either group violated normality, the two-sided Wilcoxon rank-sum test, also known as the Mann-Whitney U test, was used. Values are reported as mean ± SD unless otherwise specified. A P value < 0.05 was considered statistically significant. This statistical analysis was used to demonstrate clinical software applicability and was not intended to establish diagnostic classification performance.

### 4.1. Synthetic RR interval validation

Deterministic synthetic RR interval files were generated to test the algorithmic behavior of HRV-GUI under controlled input conditions. The validation set included constant RR sequences, alternating RR sequences, outlier-containing RR sequences, LF-dominant sinusoidal RR modulation, and HF-dominant sinusoidal RR modulation. These test patterns were generated in both human and rat RR interval ranges. Each file was loaded directly into HRV-GUI using the Load RR function, allowing the HRV computation pipeline to be evaluated independently of ECG signal quality, preprocessing, and R-peak detection. The synthetic RR validation cases and their expected analytical behavior are summarized in Table 3.

**Table 3.** Synthetic RR validation datasets and analytically expected behavior.

| **Validation case** | **Species** | **RR pattern** | **Expected analytical behavior** |
| --- | --- | --- | --- |
| Human constant | Human | 1000 ms constant RR | HR = 60 bpm; zero variability; ratio-based variables undefined |
| Human alternating | Human | Alternating 900 and 1100 ms | HR = 60 bpm; RMSSD approximately 200 ms; SD1 approximately 141.4 ms; pNN50 = 100% |
| Human outlier | Human | Mostly 1000 ms with artificial abnormal RR interval | Mean HR approximately preserved; variability metrics increase compared with constant RR |
| Human LF-dominant | Human | Sinusoidal RR modulation in human LF band | LF > HF; LF/HF > 1; LFn high and HFn low |
| Human HF-dominant | Human | Sinusoidal RR modulation in human HF band | HF > LF; LF/HF < 1; HFn high and LFn low |
| Rat constant | Rat | 174 ms constant RR | HR approximately 345 bpm; zero variability; ratio-based variables undefined |
| Rat alternating | Rat | Alternating 164 and 184 ms | HR approximately 345 bpm; RMSSD approximately 20 ms; SD1 approximately 14.1 ms; pNN50 = 0% |
| Rat outlier | Rat | Mostly 174 ms with artificial abnormal RR interval | Mean HR approximately preserved; variability metrics increase compared with constant RR |
| Rat LF-dominant | Rat | Sinusoidal RR modulation in rat LF band | LF > HF; LF/HF > 1; LFn high and HFn low |
| Rat HF-dominant | Rat | Sinusoidal RR modulation in rat HF band | HF > LF; LF/HF < 1; HFn high and LFn low |

Expected outputs were derived analytically from the known RR interval sequences. Heart rate was calculated as 60 divided by the mean RR interval in seconds. Successive RR differences were used to verify RMSSD, SDSD, and pNN50 behavior. Poincaré-derived SD1 and SD2 were calculated from consecutive RR interval pairs. For constant RR files, variability-based parameters were expected to approach zero, while SI and ratio-based spectral parameters were expected to be undefined because variability and/or spectral power were absent. For LF- and HF-dominant sinusoidal RR files, validation focused on correct frequency-band dominance rather than exact theoretical spectral power, because absolute LF and HF power values depend on interpolation, windowing, frequency resolution, and periodogram settings. The expected-versus-calculated validation results generated by HRV-GUI are summarized in Table 4.

**Table 4.**
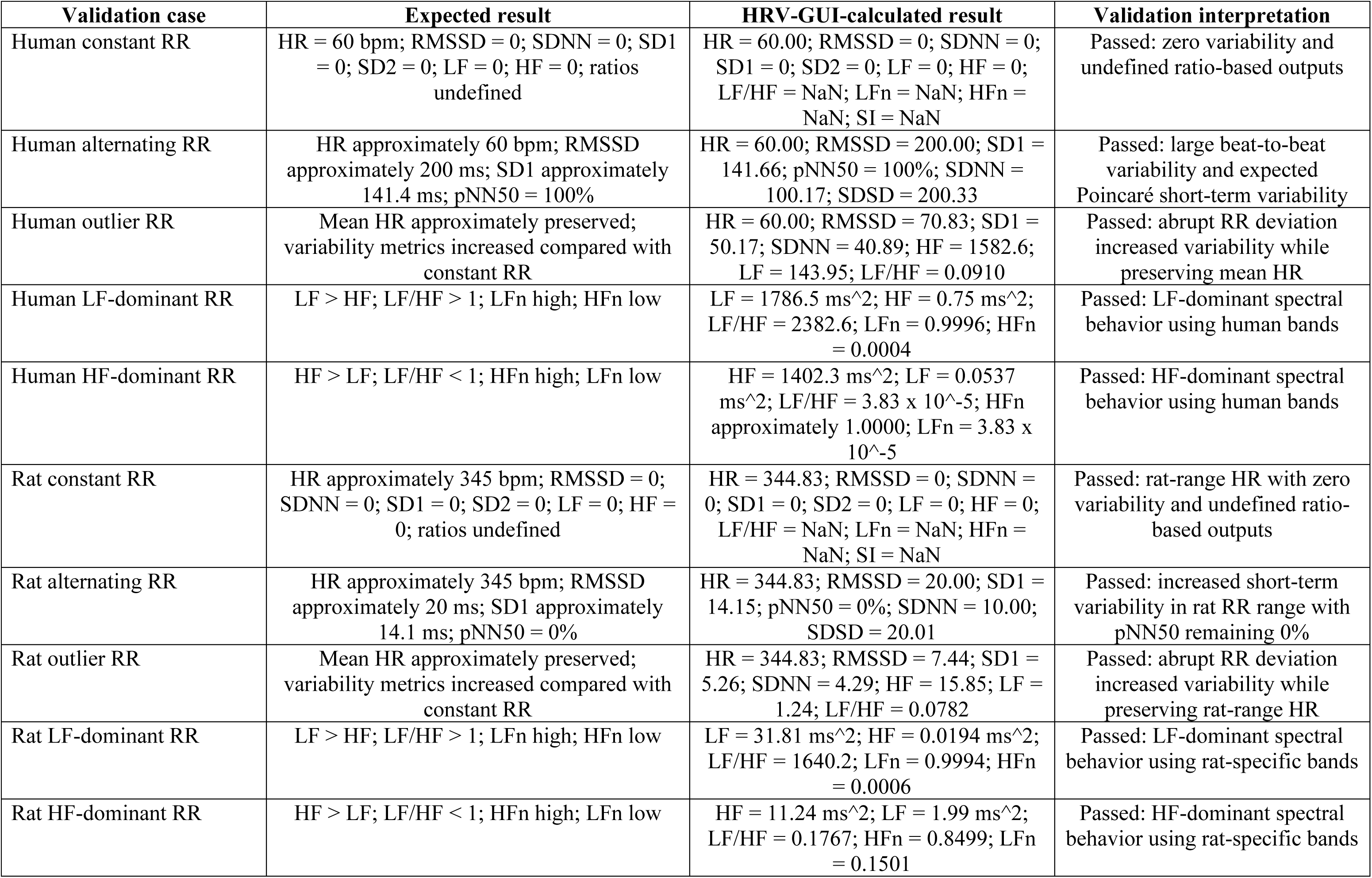
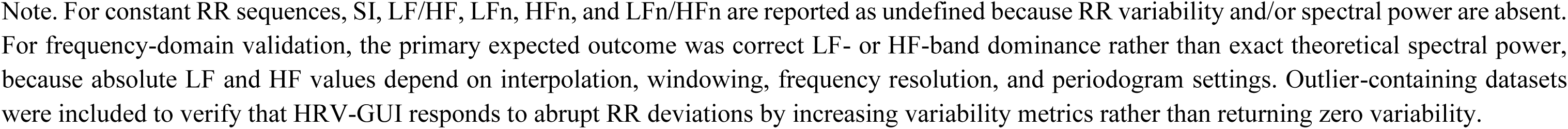
Synthetic RR validation: expected versus HRV-GUI-calculated values. Note. For constant RR sequences, SI, LF/HF, LFn, HFn, and LFn/HFn are reported as undefined because RR variability and/or spectral power are absent. For frequency-domain validation, the primary expected outcome was correct LF- or HF-band dominance rather than exact theoretical spectral power, because absolute LF and HF values depend on interpolation, windowing, frequency resolution, and periodogram settings. Outlier-containing datasets were included to verify that HRV-GUI responds to abrupt RR deviations by increasing variability metrics rather than returning zero variability.

| Validation case | Expected result | HRV-GUI-calculated result | Validation interpretation |
| --- | --- | --- | --- |
| Human constant RR | HR = 60 bpm; RMSSD = 0; SDNN = 0; SD1 = 0; SD2 = 0; LF = 0; HF = 0; ratios undefined | HR = 60.00; RMSSD = 0; SDNN = 0; SD1 = 0; SD2 = 0; LF = 0; HF = 0; LF/HF = NaN; LFn = NaN; HFn = NaN; SI = NaN | Passed: zero variability and undefined ratio-based outputs |
| Human alternating RR | HR approximately 60 bpm; RMSSD approximately 200 ms; SD1 approximately 141.4 ms; pNN50 = 100% | HR = 60.00; RMSSD = 200.00; SD1 = 141.66; pNN50 = 100%; SDNN = 100.17; SDSD = 200.33 | Passed: large beat-to-beat variability and expected Poincaré short-term variability |
| Human outlier RR | Mean HR approximately preserved; variability metrics increased compared with constant RR | HR = 60.00; RMSSD = 70.83; SD1 = 50.17; SDNN = 40.89; HF = 1582.6; LF = 143.95; LF/HF = 0.0910 | Passed: abrupt RR deviation increased variability while preserving mean HR |
| Human LF-dominant RR | LF > HF; LF/HF > 1; LFn high; HFn low | LF = 1786.5 ms<sup>2</sup>; HF = 0.75 ms<sup>2</sup>; LF/HF = 2382.6; LFn = 0.9996; HFn = 0.0004 | Passed: LF-dominant spectral behavior using human bands |
| Human HF-dominant RR | HF > LF; LF/HF < 1; HFn high; LFn low | HF = 1402.3 ms<sup>2</sup>; LF = 0.0537 ms<sup>2</sup>; LF/HF = 3.83 x 10<sup>-5</sup>; HFn approximately 1.0000; LFn = 3.83 x 10<sup>-5</sup> | Passed: HF-dominant spectral behavior using human bands |
| Rat constant RR | HR approximately 345 bpm; RMSSD = 0; SDNN = 0; SD1 = 0; SD2 = 0; LF = 0; HF = 0; ratios undefined | HR = 344.83; RMSSD = 0; SDNN = 0; SD1 = 0; SD2 = 0; LF = 0; HF = 0; LF/HF = NaN; LFn = NaN; HFn = NaN; SI = NaN | Passed: rat-range HR with zero variability and undefined ratio-based outputs |
| Rat alternating RR | HR approximately 345 bpm; RMSSD approximately 20 ms; SD1 approximately 14.1 ms; pNN50 = 0% | HR = 344.83; RMSSD = 20.00; SD1 = 14.15; pNN50 = 0%; SDNN = 10.00; SDSD = 20.01 | Passed: increased short-term variability in rat RR range with pNN50 remaining 0% |
| Rat outlier RR | Mean HR approximately preserved; variability metrics increased compared with constant RR | HR = 344.83; RMSSD = 7.44; SD1 = 5.26; SDNN = 4.29; HF = 15.85; LF = 1.24; LF/HF = 0.0782 | Passed: abrupt RR deviation increased variability while preserving rat-range HR |
| Rat LF-dominant RR | LF > HF; LF/HF > 1; LFn high; HFn low | LF = 31.81 ms<sup>2</sup>; HF = 0.0194 ms<sup>2</sup>; LF/HF = 1640.2; LFn = 0.9994; HFn = 0.0006 | Passed: LF-dominant spectral behavior using rat-specific bands |
| Rat HF-dominant RR | HF > LF; LF/HF < 1; HFn high; LFn low | HF = 11.24 ms<sup>2</sup>; LF = 1.99 ms<sup>2</sup>; LF/HF = 0.1767; HFn = 0.8499; LFn = 0.1501 | Passed: HF-dominant spectral behavior using rat-specific bands |

This validation design allowed the calculation engine to be tested under known deterministic conditions before applying the full ECG-based workflow to biological human and rat recordings.

Overall, HRV-GUI outputs matched the analytically expected behavior across all synthetic human and rat RR validation files. Constant RR files produced zero variability and undefined ratio-based variables. Alternating RR files produced the expected RMSSD, SD1, and pNN50 behavior in the human and rat ranges. LF- and HF-dominant sinusoidal RR files produced the expected spectral dominance using species-specific frequency bands. These results support the internal computational validity of the HRV-GUI calculation pipeline.

### 4.2. Cross-species baseline feasibility validation

To verify cross-species feasibility, baseline HRV outputs from healthy human controls and healthy rats were summarized descriptively in separate tables. Human results are presented in Table 5, including published comparator/reference values for selected short-term HRV parameters. Rat results are presented separately in Table 6 as descriptive feasibility outputs without a formal reference-range comparison. This separation was used to avoid implying direct statistical or biological comparison between species.

**Table 5.**
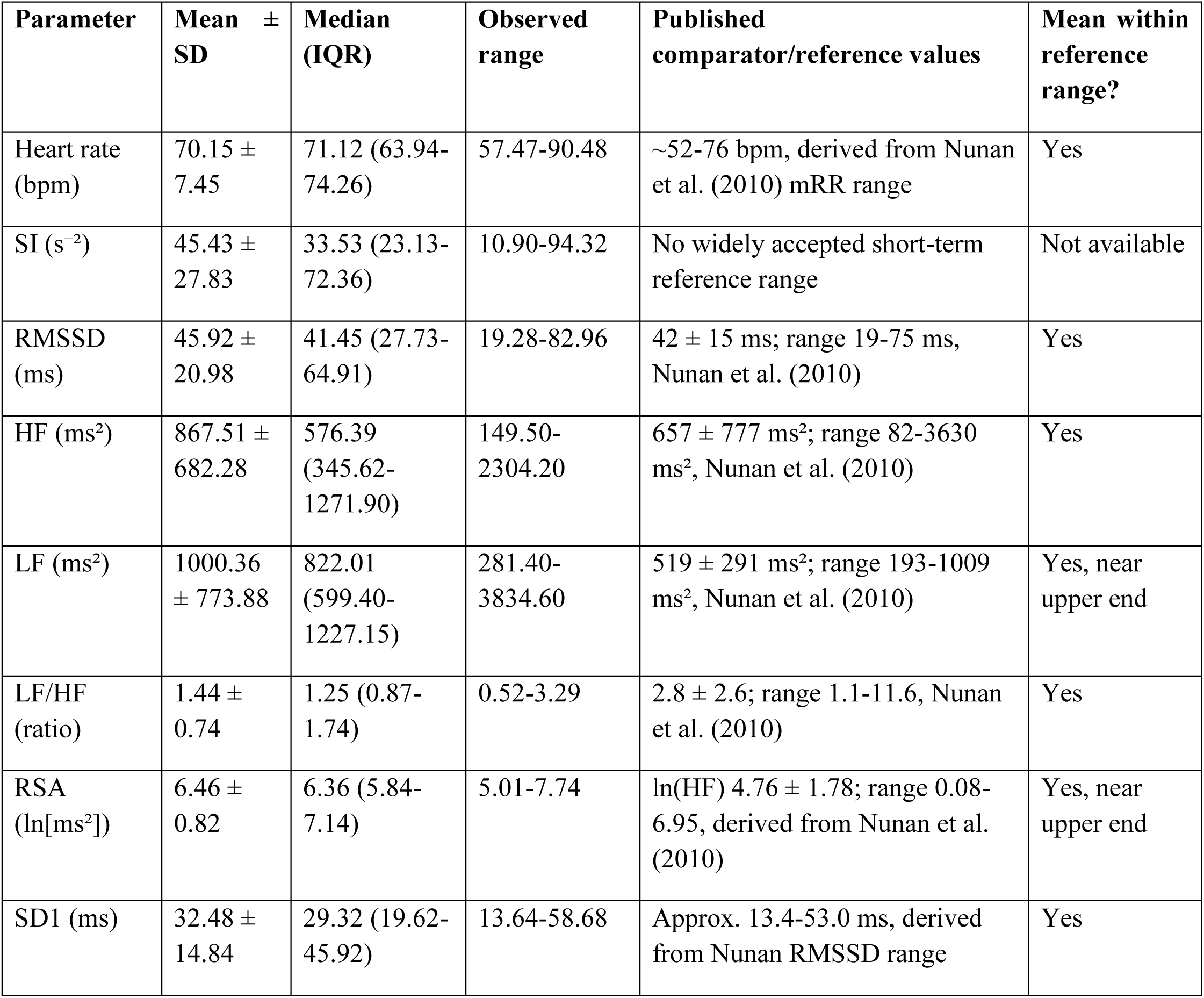
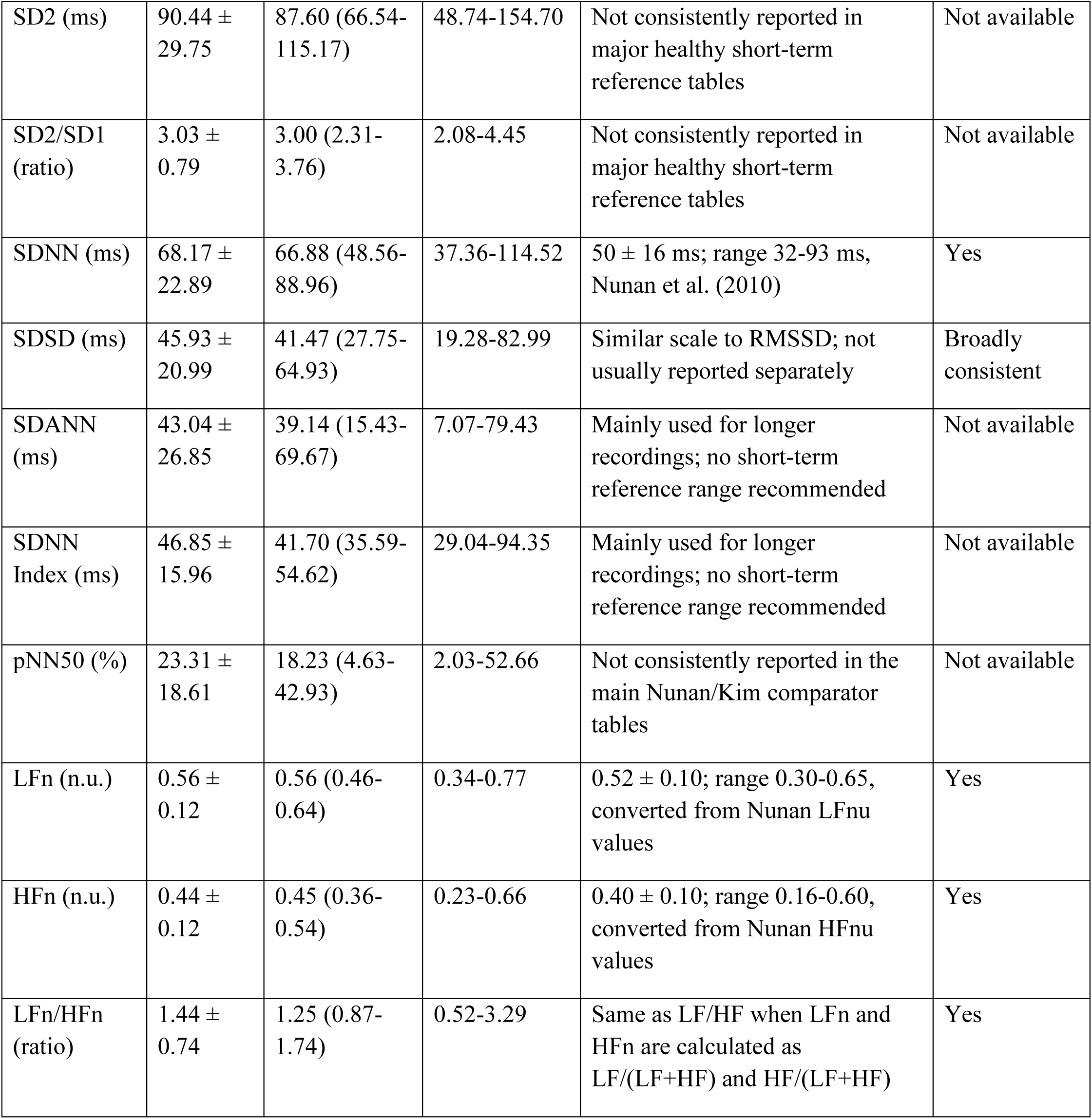
Healthy human baseline HRV outputs generated by HRV-GUI.

**Table 6.** Healthy rat HRV outputs generated by HRV-GUI.

| Parameter | Mean ± SD | Median (IQR) | Observed range |
| --- | --- | --- | --- |
| Heart rate (bpm) | 352.62 ± 31.03 | 345.12 (339.41-363.24) | 293.83-417.19 |
| SI (s<sup>-2</sup>) | 1258.58 ± 491.79 | 1167.35 (986.88-1552.05) | 407.60-2061.00 |
| RMSSD (ms) | 6.62 ± 2.40 | 6.35 (5.38-6.90) | 3.54-11.39 |
| HF (ms<sup>2</sup>) | 42.87 ± 32.09 | 35.03 (22.63-61.16) | 4.59-118.39 |
| LF (ms<sup>2</sup>) | 47.16 ± 33.84 | 45.77 (26.50-56.17) | 7.09-134.55 |
| LF/HF (ratio) | 1.21 ± 0.34 | 1.16 (1.00-1.37) | 0.79-1.87 |
| RSA (ln[ms<sup>2</sup>]) | 3.43 ± 0.96 | 3.55 (3.10-4.11) | 1.52-4.77 |
| SD1 (ms) | 4.68 ± 1.70 | 4.49 (3.81-4.88) | 2.50-8.05 |
| SD2 (ms) | 24.40 ± 9.25 | 21.94 (18.14-30.85) | 11.06-42.00 |
| SD2/SD1 (ratio) | 5.46 ± 1.85 | 5.14 (4.66-7.01) | 2.30-8.50 |
| SDNN (ms) | 17.62 ± 6.50 | 15.84 (13.78-21.99) | 8.02-30.22 |
| SDSD (ms) | 6.62 ± 2.40 | 6.35 (5.38-6.90) | 3.54-11.39 |
| SDANN (ms) | 10.77 ± 6.44 | 8.67 (6.52-12.83) | 3.49-24.64 |
| SDNN Index (ms) | 13.45 ± 4.32 | 13.06 (10.22-16.12) | 7.53-22.25 |
| pNN50 (%) | 0.23 ± 0.28 | 0.09 (0.04-0.27) | 0.00-0.88 |
| LFn (n.u.) | 0.54 ± 0.07 | 0.54 (0.50-0.58) | 0.44-0.65 |
| HFn (n.u.) | 0.46 ± 0.07 | 0.46 (0.42-0.50) | 0.35-0.56 |
| LFn/HFn (ratio) | 1.21 ± 0.34 | 1.16 (1.00-1.37) | 0.79-1.87 |

The human and rat data were not analyzed using formal inferential statistics because the two species differ substantially in heart rate, RR interval scale, autonomic physiology, respiratory rhythm, and frequency-domain definitions. Instead, the purpose of this analysis was to confirm that HRV-GUI can generate physiologically plausible outputs from both human and rodent ECG recordings using a unified workflow with species-specific settings.

For the healthy human dataset, HRV-GUI produced baseline values that were broadly consistent with published short-term HRV comparator values in healthy adults. Nunan et al. reported short-term healthy adult values including SDNN of approximately 50 ± 16 ms, RMSSD of 42 ± 15 ms, LF power of 519 ± 291 ms², HF power of 657 ± 777 ms², LFnu of 52 ± 10, HFnu of 40 ± 10, and LF/HF ratio of 2.8 ± 2.6, while also emphasizing substantial between-study variability in HRV measures (Nunan et al., 2010). Similarly, Kim and Woo reported resting 5-minute HRV values including SDNN of 39.6 ± 22.1 ms, RMSSD of 29.7 ± 18.1 ms, LF power of 417.3 ± 807.6 ms², HF power of 254.1 ± 414.1 ms², and LF/HF ratio of 2.4 ± 20.9 in a large healthy adult population (Kim and Woo, 2011). In the present healthy human dataset, HRV-GUI generated heart rate of 70.15 ± 7.45 bpm, RMSSD of 45.92 ± 20.98 ms, SDNN of 68.17 ± 22.89 ms, HF power of 867.51 ± 682.28 ms², LF power of 1000.36 ± 773.88 ms², LFn of 0.56 ± 0.12, HFn of 0.44 ± 0.12, and LF/HF ratio of 1.44 ± 0.74. As shown in Table 5, the group mean values for the main available human HRV parameters were within or broadly consistent with published comparator ranges, supporting the physiological plausibility of the human HRV-GUI outputs.

For the healthy rat dataset, HRV-GUI generated a mean heart rate of 352.62 ± 31.03 bpm, which is consistent with published values reported in healthy freely moving rats. For example, Carnevali et al. reported mean heart rates of 344.1 ± 27.1 bpm in male rats and 362.5 ± 21.9 bpm in female rats (Carnevali et al., 2023). Other rat HRV parameters were presented descriptively rather than compared against strict reference ranges because standardized normal HRV ranges for rats are not well established and are highly dependent on strain, sex, recording condition, restraint or anesthesia status, recovery period, circadian timing, segment selection, artifact handling, and frequency-band definitions. Therefore, Table 6 is intended to demonstrate that HRV-GUI can generate rodent HRV outputs across time-domain, frequency-domain, nonlinear, and Poincaré-based measures, rather than to define normal rat HRV values.

Frequency-domain outputs were successfully generated in both species using species-appropriate frequency bands. Human short-term HRV analysis commonly uses conventional human LF and HF frequency bands, whereas rodent HRV analysis requires higher frequency bands because of faster heart rate and respiratory rhythm (Task Force, 1996; Baudrie et al., 2007; Thireau et al., 2008). Because human and rat frequency-domain settings differ, the human and rat spectral outputs should be interpreted as species-specific feasibility results rather than direct biological comparisons.

Overall, the cross-species baseline results demonstrate that HRV-GUI can process ECG recordings from both clinical human and preclinical rat studies and produce physiologically plausible HR and HRV outputs. The healthy human results support agreement with published short-term HRV comparator values, while the rat results support practical feasibility of rodent ECG processing using species-specific analysis settings.

### 4.3. Clinical human use-case validation: healthy controls versus diabetic gastroparesis

To further evaluate the practical biomedical utility of HRV-GUI, we analyzed a clinical human use-case dataset consisting of 20 healthy controls and 20 patients with diabetic gastroparesis. The purpose of this analysis was not to develop a diagnostic classifier, but to test whether the complete HRV-GUI workflow could process real clinical ECG recordings and generate physiologically meaningful differences between a healthy group and a disease group.

The HC-DGP comparison was selected because gastroparesis, particularly diabetic gastroparesis, has been associated with altered autonomic regulation and reduced HRV in previous studies (Stocker et al., 2016; Nguyen et al., 2020; Ali and Chen, 2023). Therefore, the HC-DGP dataset provided a relevant clinical example for testing whether HRV-GUI could detect expected autonomic alterations across multiple HRV domains, including heart rate, stress-index-based sympathetic assessment, vagal-related measures, spectral power, time-domain variability, and Poincaré-derived indices.

For each subject, ECG recordings were processed using the HRV-GUI workflow, including ECG review, R-peak detection and correction when needed, RR interval generation, HRV computation, and result export. Summary statistics for heart rate and HRV parameters were calculated directly from the HC and DGP results. Between-group comparisons were performed as described in the Methods, and the results are summarized in Table 7. The main clinical use-case findings are also illustrated in Figure 6, which shows group differences in heart rate, SI, RMSSD, HF power, RSA, and SD1 between HC and DGP subjects. Comparisons of the remaining HRV parameters between HC and DGP subjects are shown in Supplementary Figures 1 and 2.

**Table 7.** Clinical use-case validation results comparing healthy controls and patients with diabetic gastroparesis. Values are presented as mean ± SD. HC, healthy controls; DGP, diabetic gastroparesis; HR, heart rate; SI, sympathetic index; RSA, respiratory sinus arrhythmia. Significance labels: ns, not significant; **P < 0.01; ***P < 0.001.

| Parameter | Units | HC (mean ± SD) | DGP (mean ± SD) | P value | Significance |
| --- | --- | --- | --- | --- | --- |
| HR | bpm | 70.15 ± 7.45 | 79.81 ± 12.84 | 0.0067 | ** |
| SI | s<sup>-2</sup> | 45.43 ± 27.83 | 140.65 ± 117.99 | 0.00025 | *** |
| RMSSD | ms | 45.92 ± 20.98 | 39.16 ± 60.18 | 0.0016 | ** |
| RSA | ln(ms<sup>2</sup>) | 6.46 ± 0.82 | 4.06 ± 2.03 | <0.0001 | *** |
| HF | ms<sup>2</sup> | 867.51 ± 682.28 | 348.87 ± 818.16 | <0.0001 | *** |
| LF | ms<sup>2</sup> | 1000.36 ± 773.88 | 263.20 ± 367.25 | <0.0001 | *** |
| LF/HF | ratio | 1.44 ± 0.74 | 2.23 ± 1.61 | 0.2036 | ns |
| SD1 | ms | 32.48 ± 14.84 | 27.71 ± 42.58 | 0.0016 | ** |
| SD2 | ms | 90.44 ± 29.75 | 49.77 ± 33.73 | 0.0002 | *** |
| SD2/SD1 | ratio | 3.04 ± 0.79 | 3.85 ± 3.08 | 0.8392 | ns |
| SDNN | ms | 68.17 ± 22.89 | 42.18 ± 36.21 | 0.0004 | *** |
| SDSD | ms | 45.93 ± 20.99 | 39.18 ± 60.20 | 0.0016 | ** |
| SDANN | ms | 43.04 ± 26.85 | 11.48 ± 6.39 | <0.0001 | *** |
| SDNN Index | ms | 46.85 ± 15.96 | 37.88 ± 35.53 | 0.0013 | ** |
| pNN50 | % | 23.31 ± 18.61 | 11.38 ± 25.50 | 0.0006 | *** |
| LFn | n.u | 0.56 ± 0.12 | 0.62 ± 0.17 | 0.2094 | ns |
| HFn | n.u | 0.44 ± 0.12 | 0.38 ± 0.17 | 0.2094 | ns |
| LFn/HFn | ratio | 1.44 ± 0.74 | 2.23 ± 1.61 | 0.2036 | ns |

**Figure 6.**
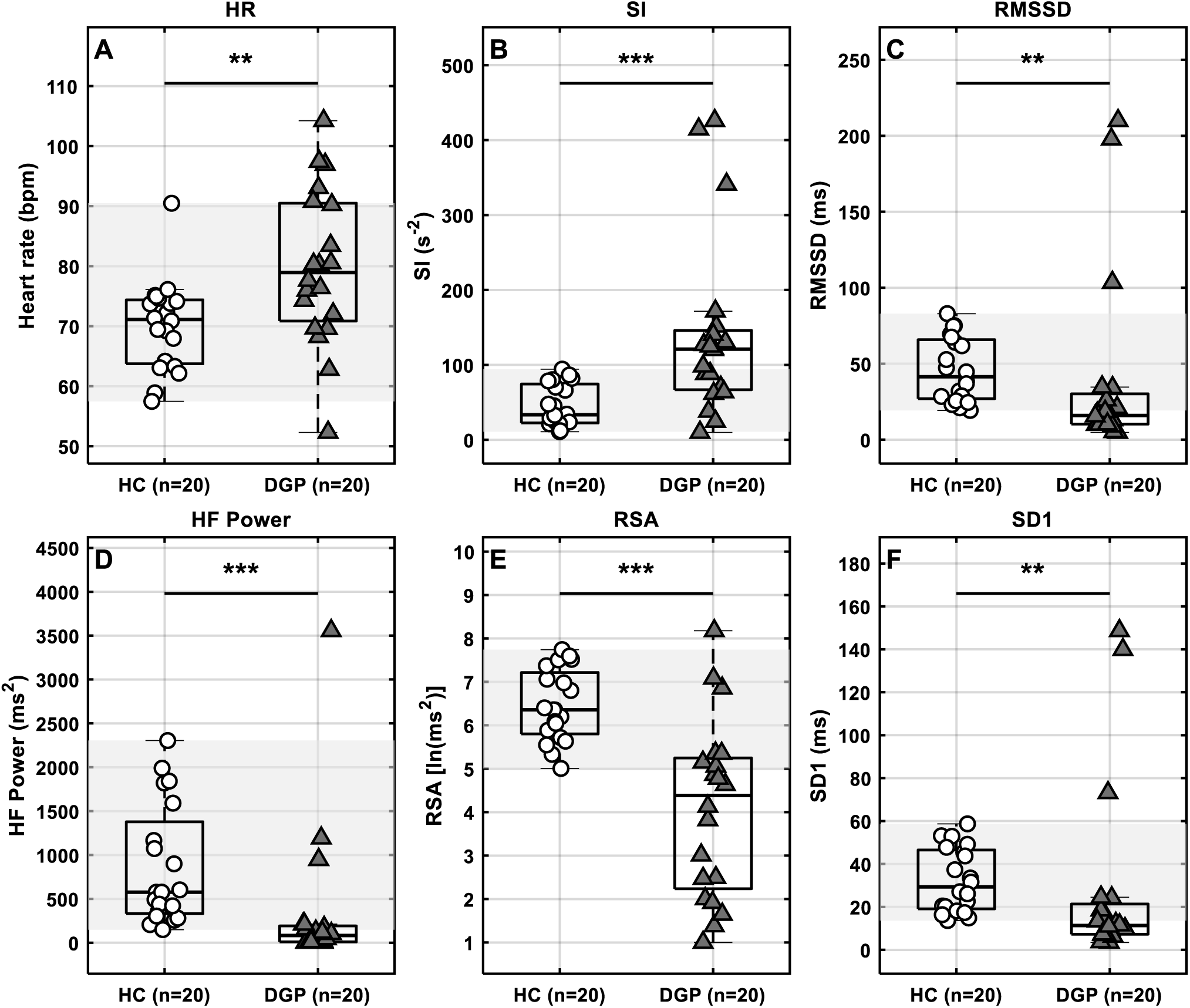
HRV-GUI clinical use-case validation in healthy controls and diabetic gastroparesis. HRV-GUI outputs were compared between healthy controls (HC, n = 20) and patients with diabetic gastroparesis (DGP, n = 20). The figure shows (A) heart rate, (B) Sympathetic Index (SI), (C) RMSSD, (D) HF power, (E) RSA, and (F) SD1. Open circles represent HC subjects, and filled triangles represent DGP patients. Significance labels are shown above each subpanel. The gray shaded region represents the empirical 2.5th–97.5th percentile interval of the HC group.

The HC-DGP comparison showed several statistically significant between-group differences. Heart rate was higher in DGP than in HC subjects (79.81 ± 12.84 vs. 70.15 ± 7.45 bpm, P = 0.0067). SI was also substantially higher in DGP than in HC subjects (140.65 ± 117.99 vs. 45.43 ± 27.83 s⁻², P = 0.00025), consistent with greater stress-index-based sympathetic predominance or reduced autonomic flexibility (Baevsky and Berseneva, 2008). Because SI is a geometric HRV-derived stress index rather than a direct sympathetic nerve recording, this finding was interpreted as an HRV-based marker of altered autonomic regulation.

In contrast, several vagal and overall HRV-related measures were lower in DGP than in HC subjects. RMSSD was lower in DGP than in HC subjects (39.16 ± 60.18 vs. 45.92 ± 20.98 ms, P = 0.0016), and SD1 showed the same pattern (27.71 ± 42.58 vs. 32.48 ± 14.84 ms, P = 0.0016). Because SD1 is closely related to short-term beat-to-beat variability and RMSSD, these parallel findings support reduced short-term HRV in the DGP group (Shaffer and Ginsberg, 2017). RSA was also markedly lower in DGP than in HC subjects (4.06 ± 2.03 vs. 6.46 ± 0.82 ln[ms²], P < 0.0001), and HF power was lower in DGP than in HC subjects (348.87 ± 818.16 vs. 867.51 ± 682.28 ms², P < 0.0001). Since HF power and RSA are commonly interpreted as indices related to respiratory-linked vagal modulation, these findings suggest reduced vagal-related HRV in DGP (Task Force, 1996; Shaffer and Ginsberg, 2017).

Additional significant differences were observed in LF power, SD2, SDNN, SDSD, SDANN, SDNN Index, and pNN50. LF power was lower in DGP than in HC subjects (263.20 ± 367.25 vs. 1000.36 ± 773.88 ms², P < 0.0001), SD2 was lower in DGP (49.77 ± 33.73 vs. 90.44 ± 29.75 ms, P = 0.0002), SDNN was lower in DGP (42.18 ± 36.21 vs. 68.17 ± 22.89 ms, P = 0.0004), SDSD was lower in DGP (39.17 ± 60.20 vs. 45.93 ± 20.99 ms, P = 0.0016), SDANN was markedly lower in DGP (11.48 ± 6.39 vs. 43.04 ± 26.85 ms, P < 0.0001), SDNN Index was lower in DGP (37.88 ± 35.53 vs. 46.85 ± 15.96 ms, P = 0.0013), and pNN50 was lower in DGP (11.38 ± 25.50 vs. 23.31 ± 18.61%, P = 0.0006). Together, these findings support a broader reduction in absolute HRV magnitude in DGP.

The observed pattern of higher heart rate and SI together with lower RSA, HF power, RMSSD, SD1, SD2, SDNN, SDSD, SDANN, SDNN Index, LF power, and pNN50 in DGP is consistent with altered autonomic regulation and reduced overall HRV reported in gastroparesis and related gastrointestinal disorders (Mazurak et al., 2012; Stocker et al., 2016; Nguyen et al., 2020; Ali and Chen, 2023). Overall, this clinical use-case analysis supports the ability of the HRV-GUI software to generate clinically interpretable autonomic differences in diabetic gastroparesis.

However, normalized spectral indices were not significantly different between groups. LF/HF, LFn, HFn, and LFn/HFn did not differ significantly between HC and DGP subjects. This suggests that the clinical use-case dataset primarily demonstrated reduced absolute HRV power and reduced vagal/overall variability rather than a consistent shift in normalized LF-HF balance. This distinction is important because absolute LF and HF powers can decrease substantially while normalized LF-HF proportions remain statistically similar.

## 5. Discussion and significance

This manuscript presents HRV-GUI, a MATLAB-based graphical user interface for ECG-derived HRV analysis in human and rodent biomedical research. The software was designed as a workflow-oriented research interface rather than a simple HRV calculator. A central contribution of this study is that HRV-GUI was evaluated not only as a software interface, but also as a biomedical HRV analysis workflow validated using controlled synthetic RR files, healthy human ECG recordings, healthy rat ECG recordings, and a clinical HC-DGP comparison. It allows users to move from ECG waveform inspection to finalized HRV outputs through a transparent sequence of steps while preserving user control over the analysis decisions that most strongly influence HRV results.

A key advantage of HRV-GUI is that it maintains user oversight during ECG segment selection, R-peak detection thresholding, peak-distance adjustment, manual peak correction, RR interval review, diagnostic visualization, and final result export. This is important because HRV metrics are highly sensitive to missed beats, false peaks, noise, motion artifact, ectopic beats, and inappropriate segment selection. These issues have been recognized in HRV standards and artifact-processing approaches (Task Force, 1996; Kaufmann et al., 2011). By allowing direct visual inspection of both ECG waveforms and RR interval series, HRV-GUI helps users identify potential signal-quality or peak-detection problems before final autonomic indices are calculated.

The inclusion of both human and rat processing is a practical strength of the software. Many HRV tools are optimized primarily for human recordings, whereas translational laboratories often require consistent workflow for both clinical and preclinical ECG datasets. HRV-GUI addresses this need by maintaining a unified interface while allowing species-specific settings for peak detection and frequency-domain analysis. The cross-species feasibility results showed that the software could process real human and rodent ECG recordings and generate physiologically plausible outputs. In healthy humans, the HRV-GUI outputs were broadly consistent with published short-term HRV comparator values (Nunan et al., 2010; Kim and Woo, 2011). In rats, the software generated descriptive HRV outputs across multiple HRV domains, with heart rate values consistent with published healthy rat values, while recognizing that standardized rat HRV reference ranges are not well established and are strongly influenced by strain, sex, recording condition, restraint or anesthesia status, recovery period, circadian timing, segment selection, artifact handling, and frequency-band definitions (Baudrie et al., 2007; Thireau et al., 2008; Carnevali et al., 2023).

Synthetic RR validation provided an important computational test that was independent of ECG signal quality and R-peak detection performance. Constant, alternating, outlier-containing, LF-dominant, and HF-dominant RR files allowed the calculation engine to be tested under deterministic conditions where the expected behavior was known in advance. The agreement between the expected analytical behavior and HRV-GUI outputs supports the internal validity of the implemented HRV calculations.

The HC-DGP clinical use-case analysis further demonstrated that the HRV-GUI could generate clinically interpretable outputs from real human recordings. This analysis was not designed to establish a definitive diagnostic classifier for diabetic gastroparesis; instead, it was used to show that the software workflow can detect biologically meaningful autonomic differences between a healthy control group and a disease group. In this dataset, patients with diabetic gastroparesis showed higher heart rate and SI, together with lower RSA, HF power, RMSSD, SD1, SD2, SDNN, SDSD, SDANN, SDNN Index, LF power, and pNN50. This pattern supports reduced overall HRV and altered autonomic regulation in DGP. These findings are consistent with broader literature showing autonomic dysfunction in gastrointestinal disorders and diabetic gastroparesis (Mazurak et al., 2012; Ali and Chen, 2023; Nguyen et al., 2020). Importantly, normalized spectral indices, including LFn, HFn, LFn/HFn, and LF/HF, were not significantly different between groups, suggesting that the clinical use-case primarily demonstrated reduced absolute HRV power rather than a consistent shift in normalized LF-HF balance.

Together, these analyses show that HRV-GUI performs as expected on controlled synthetic RR files and can be applied to real human and rat ECG recordings. The clinical HC-DGP use-case further demonstrates that the software can generate physiologically meaningful autonomic differences in a disease-relevant human dataset. These findings support HRV-GUI as a validated biomedical software workflow for HRV analysis in both clinical human studies and preclinical rodent research.

Overall, the HRV-GUI presented in this study provides a transparent and flexible MATLAB-based platform for ECG-derived HRV analysis across human and rodent biomedical research. By combining visual ECG inspection, user-guided peak correction, cross-species settings, multi-domain HRV outputs, diagnostic visualization, and export-ready results, the software offers a practical tool for laboratories studying autonomic regulation in both clinical and preclinical settings.

## 6. Limitations and future work

The present study has several scope-related limitations. First, synthetic RR validation was used to test the internal consistency of the HRV calculation engine under deterministic conditions. Although this supports the expected behavior of the implemented calculations, future studies may further compare HRV-GUI outputs with established HRV software packages using shared ECG and RR interval datasets.

Second, the HC-DGP dataset was included as a clinical use-case demonstration rather than a diagnostic biomarker study. Therefore, the clinical results should be interpreted as evidence that HRV-GUI can generate physiologically interpretable group differences in real human recordings, not as a definitive diagnostic classifier for diabetic gastroparesis. Larger studies with covariate adjustment and external validation would be needed for diagnostic biomarker development. Interpretation of individual HRV metrics should also remain cautious because commonly used vagal-related measures such as RMSSD can be influenced by respiratory conditions and may not always reflect parasympathetic reactivity in a simple one-to-one manner (Ali, Liu, et al., 2023). This is particularly relevant for clinical and translational HRV studies, where breathing pattern, recording condition, and disease status may influence HRV outputs.

Third, rat HRV outputs were presented descriptively because standardized normal HRV reference ranges for rodents are not universally established and are strongly influenced by strain, sex, recording condition, restraint or anesthesia status, recovery period, circadian timing, segment selection, artifact handling, and frequency-band definitions. Therefore, the rat analysis was framed as cross-species feasibility rather than formal reference-range validation.

Finally, HRV-GUI is presented as a research-oriented MATLAB tool. Future versions may include expanded documentation, example datasets, broader software packaging, and additional validation across different ECG acquisition systems, disease models, recording conditions, and species.

## 7. Conclusions

The HRV-GUI is a MATLAB-based graphical user interface for ECG-derived heart rate variability analysis in human and rodent biomedical research. The software integrates signal loading, preprocessing, R-peak detection and correction, RR interval generation, multi-domain HRV computation, diagnostic plotting, data export, and session handling in a transparent workflow. The validation structure demonstrated computational plausibility using synthetic RR datasets, cross-species feasibility using healthy human and rat recordings, and clinical applicability using a healthy control versus diabetic gastroparesis use-case dataset. Together, these findings support HRV-GUI as a practical and extensible tool for translational autonomic research in both clinical and preclinical settings.

## Declarations

### Conflict of interest

The authors declare no conflict of interest.

### Funding

This work was partially supported by NIH grants (UG3NS115108, 1R44AT011380)

### Ethics statement

Human ECG recordings and animal experiments were conducted under protocols approved by the University of Michigan Institutional Review Board and the University of Michigan Institutional Animal Care and Use Committee, respectively. Written informed consent was obtained from human participants as required by the approved protocols. All procedures were performed in accordance with institutional guidelines and applicable regulations.

### Data availability

The data supporting this manuscript may be made available upon reasonable request, subject to institutional and ethical approvals.

### Code availability

The HRV-GUI is a MATLAB program. The final distribution format may include source code, a compiled standalone MATLAB application, or both, subject to institutional review.

### Declaration of generative AI use

Generative AI assistance was used to help create the conceptual workflow diagram shown in Figure 1. The authors reviewed and edited the figure and take full responsibility for its content.

### Author contributions

M. Khawar Ali: conceptualization, software development, validation, formal analysis, visualization, writing-original draft, and revision. Jiande D. Z. Chen: supervision, project administration, interpretation, and manuscript review.

---

**Supplementary Figure 1.**
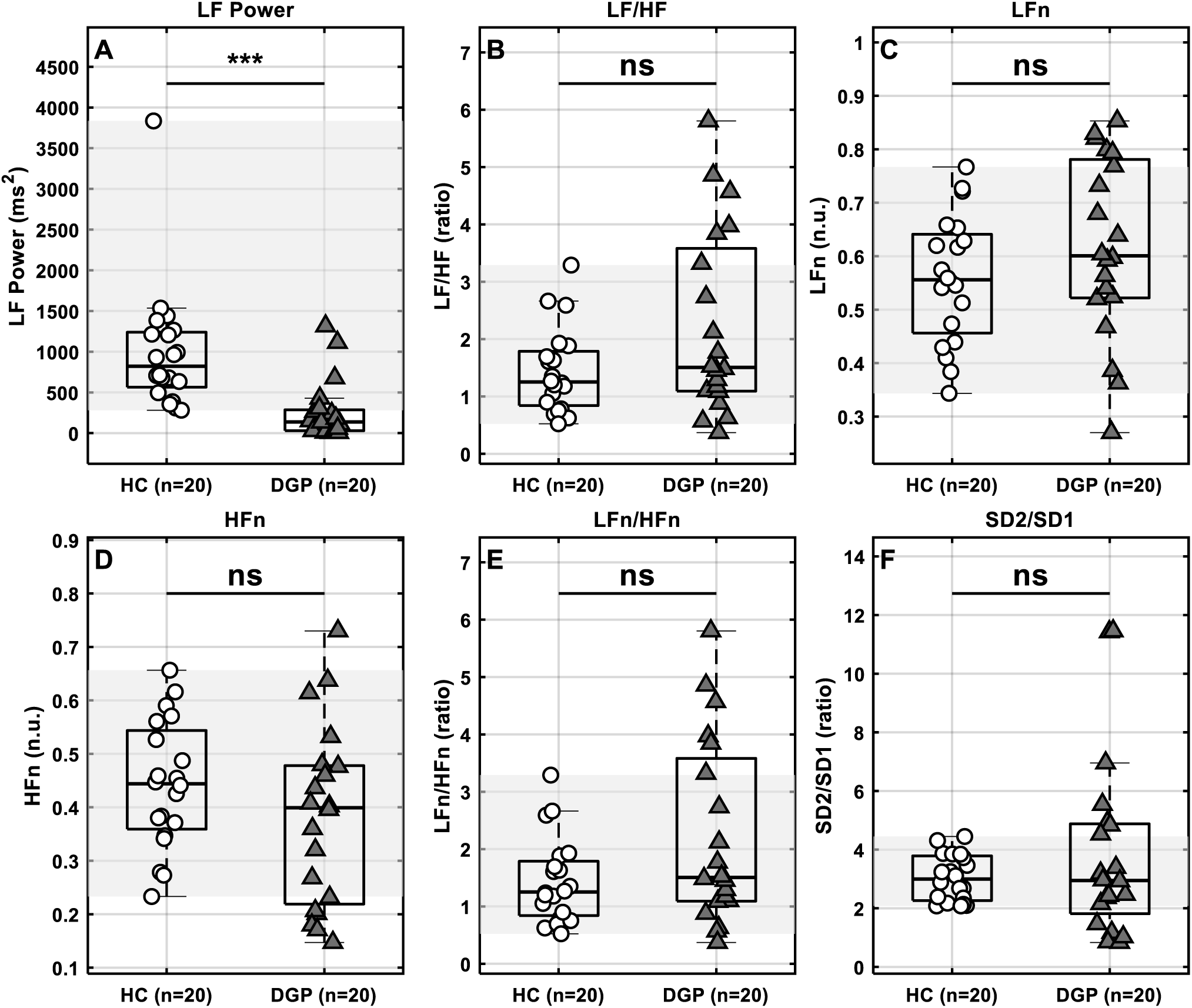
Additional HRV-GUI clinical use-case outputs in healthy controls and diabetic gastroparesis. HRV-GUI outputs were compared between healthy controls (HC, n = 20) and patients with diabetic gastroparesis (DGP, n = 20). The figure shows (A) LF power, (B) LF/HF ratio, (C) LFn, (D) HFn, (E) LFn/HFn, and (F) SD2/SD1. Open circles represent HC subjects, and filled triangles represent DGP patients. Significance labels are shown above each subpanel. The gray shaded region represents the empirical 2.5th–97.5th percentile interval of the HC group.

**Supplementary Figure 2.**
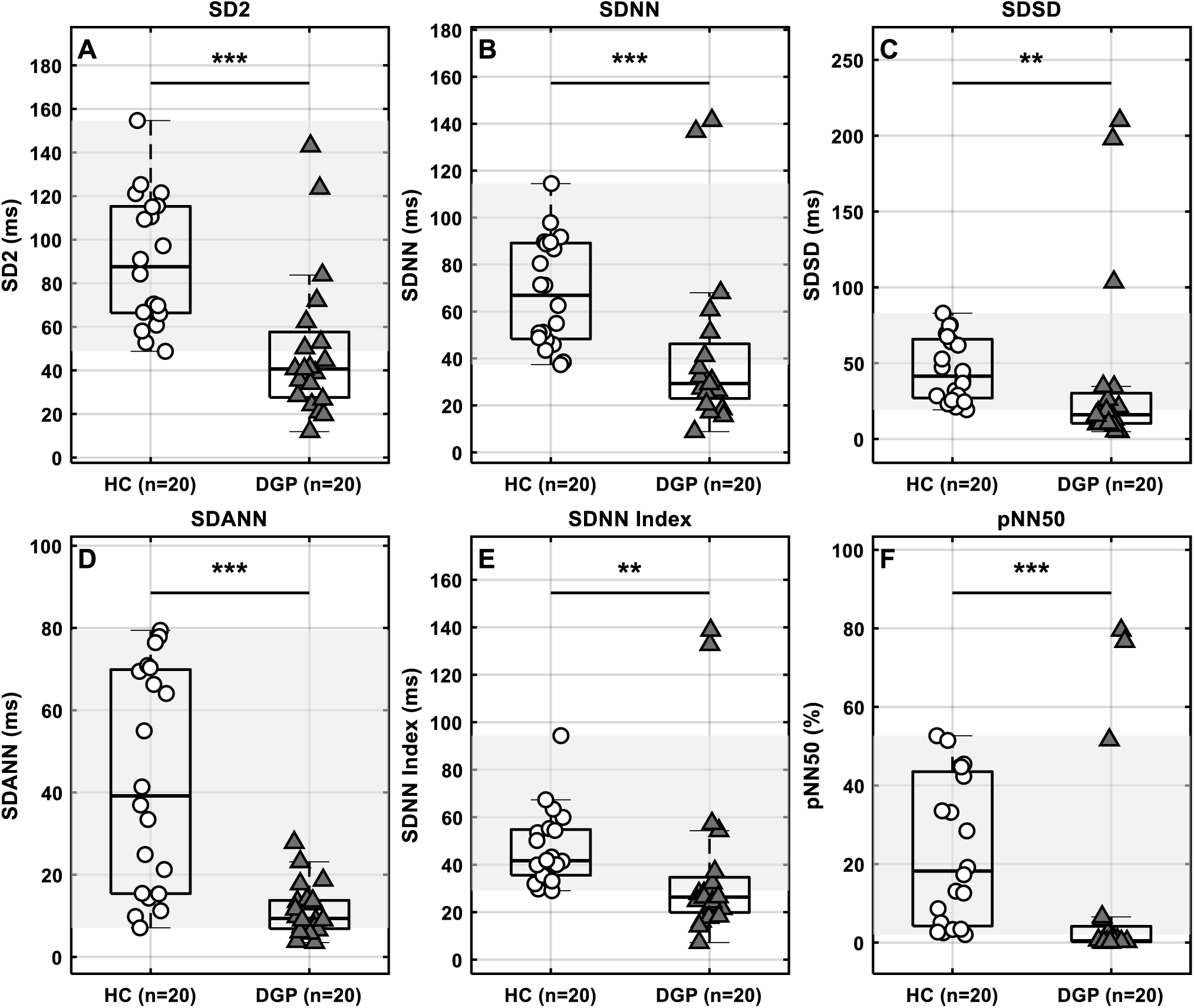
Additional HRV-GUI clinical use-case outputs in healthy controls and diabetic gastroparesis. HRV-GUI outputs were compared between healthy controls (HC, n = 20) and patients with diabetic gastroparesis (DGP, n = 20). The figure shows (A) SD2, (B) SDNN, (C) SDSD, (D) SDANN, (E) SDNN Index, and (F) pNN50%. Open circles represent HC subjects, and filled triangles represent DGP patients. Significance labels are shown above each subpanel. The gray shaded region represents the empirical 2.5th–97.5th percentile interval of the HC group.

## Notes

### Competing Interest Statement

The authors have declared no competing interest.

## References

1. Ali MK, Chen JDZ. Roles of heart rate variability in assessing autonomic nervous system in functional gastrointestinal disorders: a systematic review. Diagnostics. 2023;13(2):293. doi: 10.3390/diagnostics13020293

2. Ali MK, Gong S, Nojkov B, Burnett C, Chen JDZ. Best parameters of heart rate variability for assessing autonomic responses to brief rectal distention in patients with irritable bowel syndrome. Sensors. 2023;23(19):8128. doi: 10.3390/s23198128

3. Ali MK, Liu L, Chen JH, Huizinga JD. Optimizing autonomic function analysis via heart rate variability associated with motor activity of the human colon. Front Physiol. 2021;12:619722. doi: 10.3389/fphys.2021.619722

4. Ali MK, Liu L, Hussain A, Zheng D, Alam M, Chen JH, Huizinga JD. Root mean square of successive differences is not a valid measure of parasympathetic reactivity during slow deep breathing. Am J Physiol Regul Integr Comp Physiol. 2023;324(4):R446–R456. doi:10.1152/ajpregu.00272.2022.

5. Baevsky RM, Berseneva AP. Introduction to Prenosological Diagnostics. Moscow: Slovo; 2008.

6. Baudrie V, Laude D, Elghozi JL. Optimal frequency ranges for extracting information on cardiovascular autonomic control from the blood pressure and pulse interval spectrograms in mice. Am J Physiol Regul Integr Comp Physiol. 2007;292(2):R904–R912. doi: 10.1152/ajpregu.00488.2006

7. Brennan M, Palaniswami M, Kamen P. Do existing measures of Poincare plot geometry reflect nonlinear features of heart rate variability? IEEE Trans Biomed Eng. 2001;48(11):1342–1347. doi: 10.1109/10.959330

8. Carnevali L, Barbetti M, Statello R, Williams DP, Thayer JF, Sgoifo A. Sex differences in heart rate and heart rate variability in rats: implications for translational research. Front Physiol. 2023;14:1170320. doi: 10.3389/fphys.2023.1170320.

9. Garcia CA. A MATLAB toolbox for the analysis of heart rate variability. Conf Proc IEEE Eng Med Biol Soc. 2009;2009:3823–3826.

10. Goldberger AL, Amaral LAN, Glass L, Hausdorff JM, Ivanov PC, Mark RG, Mietus JE, Moody GB, Peng CK, Stanley HE. PhysioBank, PhysioToolkit, and PhysioNet: components of a new research resource for complex physiologic signals. Circulation. 2000;101(23):e215–e220. doi: 10.1161/01.CIR.101.23.e215

11. Kaufmann T, Sütterlin S, Schulz SM, Vögele C. ARTiiFACT: a tool for heart rate artifact processing and heart rate variability analysis. Behav Res Methods. 2011;43(4):1161–1170. doi: 10.3758/s13428-011-0107-7

12. Kim GM, Woo JM. Determinants for heart rate variability in a normal Korean population. J Korean Med Sci. 2011;26(10):1293–1298. doi: 10.3346/jkms.2011.26.10.1293.

13. Liu L, Milkova N, Nirmalathasan S, Ali MK, Sharma K, Huizinga JD, Chen JH. Diagnosis of colonic dysmotility associated with autonomic dysfunction in patients with chronic refractory constipation. Sci Rep. 2022;12:12051. doi: 10.1038/s41598-022-15945-6

14. Mazurak N, Seredyuk N, Sauer H, Teufel M, Enck P. Heart rate variability in the irritable bowel syndrome: a review of the literature. Neurogastroenterol Motil. 2012;24(3):206–216. doi: 10.1111/j.1365-2982.2011.01866.x.

15. Nguyen L, Wilson LA, Miriel L, Pasricha PJ, Kuo B, Hasler WL, McCallum RW, Sarosiek I, Koch KL, Snape WJ, et al. Autonomic function in gastroparesis and chronic unexplained nausea and vomiting: relationship with etiology, gastric emptying, and symptom severity. Neurogastroenterol Motil. 2020;32(8):e13810. doi: 10.1111/nmo.13810

16. Nunan D, Sandercock GRH, Brodie DA. A quantitative systematic review of normal values for short-term heart rate variability in healthy adults. Pacing Clin Electrophysiol. 2010;33(11):1407–1417. doi: 10.1111/j.1540-8159.2010.02841.x.

17. Shaffer F, Ginsberg JP. An overview of heart rate variability metrics and norms. Front Public Health. 2017;5:258. doi: 10.3389/fpubh.2017.00258

18. Stocker A, Abell TL, Rashed H, Kedar A, Boatright B, Chen J. Autonomic evaluation of patients with gastroparesis and neurostimulation: comparisons of direct/systemic and indirect/cardiac measures. Gastroenterol Res. 2016;9(1):10–16. doi: 10.14740/gr667w

19. Tarvainen MP, Niskanen JP, Lipponen JA, Ranta-aho PO, Karjalainen PA. Kubios HRV: heart rate variability analysis software. Comput Methods Programs Biomed. 2014;113(1):210–220. doi: 10.1016/j.cmpb.2013.07.024

20. Task Force of the European Society of Cardiology and the North American Society of Pacing and Electrophysiology. Heart rate variability: standards of measurement, physiological interpretation and clinical use. Circulation. 1996;93(5):1043–1065. doi: 10.1161/01.CIR.93.5.1043.

21. Thayer JF, Sternberg EM. Beyond heart rate variability: vagal regulation of allostatic systems. Ann N Y Acad Sci. 2006;1088:361–372. doi: 10.1196/annals.1366.014.

22. Thireau J, Zhang BL, Poisson D, Babuty D. Heart rate variability in mice: a theoretical and practical guide. Exp Physiol. 2008;93(1):83–94. doi: 10.1113/expphysiol.2007.040733

23. Vidaurre C, Sander TH, Schlögl A. BioSig: the free and open source software library for biomedical signal processing. Comput Intell Neurosci. 2011;2011:935364. doi: 10.1155/2011/935364.

24. Wang X, Yang B, Yin J, Wei W, Chen JDZ. Electroacupuncture via chronically implanted electrodes improves gastrointestinal motility by balancing sympathovagal activities in a rat model of constipation. Am J Physiol Gastrointest Liver Physiol. 2019;316(6):G797–G805. doi: 10.1152/ajpgi.00018.2018

